# Parallelized Acquisition of Orbitrap and Astral Analyzers Enables High-Throughput Quantitative Analysis

**DOI:** 10.1101/2023.06.02.543408

**Authors:** Hamish Stewart, Dmitry Grinfeld, Anastassios Giannakopulos, Johannes Petzoldt, Toby Shanley, Matthew Garland, Eduard Denisov, Amelia Peterson, Eugen Damoc, Martin Zeller, Tabiwang N. Arrey, Anna Pashkova, Santosh Renuse, Amirmansoor Hakimi, Andreas Kühn, Matthias Biel, Arne Kreutzmann, Bernd Hagedorn, Immo Colonius, Adrian Schütz, Arne Stefes, Ankit Dwivedi, Daniel Mourad, Max Hoek, Bastian Reitemeier, Philipp Cochems, Alexander Kholomeev, Robert Ostermann, Gregor Quiring, Maximilian Ochmann, Sascha Möhring, Alexander Wagner, André Petker, Sebastian Kanngiesser, Michael Wiedemeyer, Wilko Balschun, Daniel Hermanson, Vlad Zabrouskov, Alexander Makarov, Christian Hock

## Abstract

The growing trend towards high-throughput proteomics demands rapid liquid chromatography-mass spectrometry (LC-MS) cycles that limit the available time to gather the large numbers of MS/MS fragmentation spectra required for identification. Orbitrap analyzers scale performance with acquisition time, and necessarily sacrifice sensitivity and resolving power to deliver higher acquisition rates. We developed a new mass spectrometer that combines a mass resolving quadrupole, the Orbitrap and the novel Asymmetric Track Lossless (Astral) analyzer. The new hybrid instrument enables faster acquisition of high-resolution accurate mass (HRAM) MS/MS spectra compared to state-of-the-art mass spectrometers. Accordingly, new proteomics methods were developed that leverage the strengths of each HRAM analyzer, whereby the Orbitrap analyzer performs full scans with high dynamic range and resolution, synchronized with Astral analyzer’s acquisition of fast and sensitive HRAM MS/MS scans. Substantial improvements are demonstrated over previous methods using current state-of-the-art mass spectrometers.

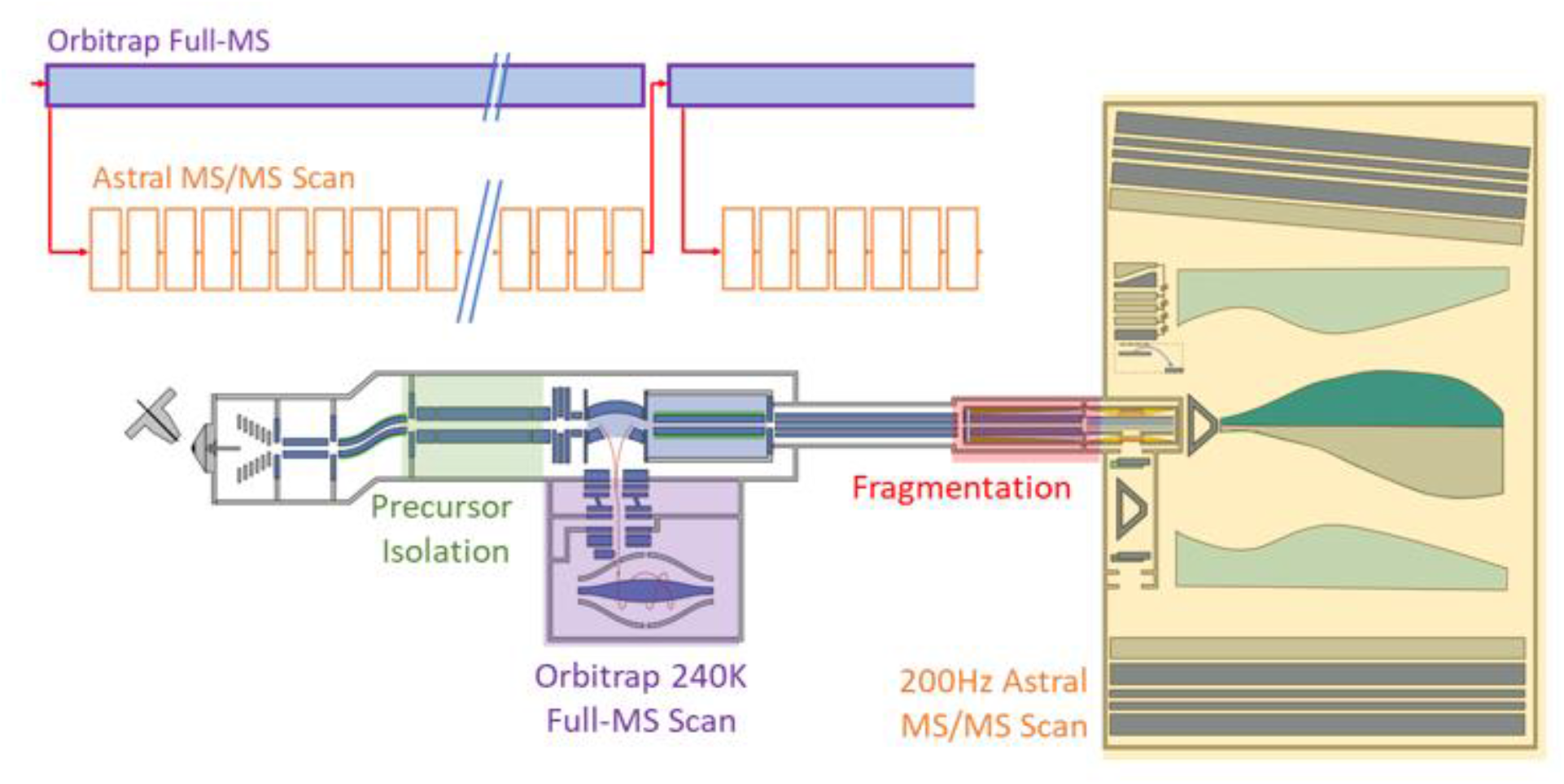

## Introduction

Bottom-up proteomics workflows profile complex mixtures of proteins through the analysis of enzymatically digested peptides delivered to a mass spectrometer via chromatographic separation^1^. In a popular data dependent workflow, the peptide precursor ions are detected within a full MS scan, isolated, fragmented, and the resulting MS/MS spectra analyzed by bioinformatic tools to identify parent peptides and infer pre-digest proteins^2-5^. A typical sample may easily contain 100,000s of peptide features^6^, with the depth of coverage defined by the mass spectrometer’s ability to generate sufficient quality spectra across a high dynamic range in the available time.

For MS/MS spectra, sensitivity has historically been the most pressing challenges, due to the extremely low abundance of many peptide features, caused by the enormous range of protein abundances^7,8^. Thermo Scientific™ Orbitrap™ mass spectrometers are important proteomics instruments due to their relatively high transmission and accurate precursor mass measurements with high dynamic range^9-12^. Pairing of an Orbitrap analyzer with a more sensitive, albeit lower resolving power, linear ion trap with single ion detection has long been an effective strategy, allowing the Orbitrap analyzer to carry out highly resolved full mass scans whilst the linear ion trap performed isolation and sensitive MS/MS acquisition^10^. An important evolution of this strategy was the Tribrid instrument architecture combining an Orbitrap analyzer for precursor detection, quadrupole mass filter for precursor isolation, and a dual-pressure linear ion trap for sensitive MS/MS^9^. A parallelized acquisition method allows each analyzer to carry out fit-for-purpose acquisition and operate efficiently across the entire instrument.

For ultra-high-throughput proteomics with chromatographic gradients compressed down to 5 or 8 minutes, large numbers of sample ions pass through the instrument within a short timeframe, and the requisite number of MS/MS spectra to identify peptides need to be generated much more quickly. An analyzer acquiring 25 spectra per second (25 Hz) to target all detectable peptides in 100 minutes, as had been estimated^6^, would have to operate above 300 Hz to manage the same MS/MS coverage in 8 minutes. Although there are ways to mitigate this fundamental burden, such as via parallel identification of multiple co-isolated precursor ions within an individual MS/MS spectrum^13-14^, there are multiple technological innovations required to reach such a challenging acquisition rate with sufficient sensitivity and dynamic range.

Orbitrap MS/MS acquisition has been limited to above 40Hz^15^, and this comes with a trade-off of sensitivity and resolution due to shorter ion detection transient. Thus, Orbitrap MS/MS methods must balance repetition rate with resolution and sensitivity^16^. Moderately faster acquisition rates of 70 Hz MS/MS are fundamentally possible with Orbitrap analyzers and demonstrates some benefit for high throughput workflows^17^.

Time-of-flight (ToF) analyzers on the other hand have typically very short ion measurement periods, tens of microseconds to a few milliseconds for ions to traverse the flight path, depending on size and design, and may routinely generate spectra at >100Hz^18,19^. Such analyzers have historically suffered from poor sensitivity, due largely to low duty cycle and transmission losses within the orthogonal accelerators that introduce ions into the analyzer^20^ and on multiple grids along the ion path. Some of these losses have been reduced with an improved accelerator design^21^ or by replacement of the device with a direct-extraction ion trap^22,23^ in a similar manner to how a curved linear trap (C-trap) is used to inject ions into Orbitrap analyzers^24^. However, severe temporal broadening of ion packets and a strong dependence on the numbers of ions combined with complex electronics and detector saturation precluded the widespread utilization of this curved liner trap approach with continuous ion beams.

Multi-reflection or multi-turn time-of-flight analyzer designs with folded and grid-free flight paths offer far higher resolving power and consequently higher statistical mass accuracy than conventional linear or single reflection designs^25^. Numerous “MR-ToF”, or “Open Electrostatic Trap” designs of this type have been introduced over decades^25-27^. In spite of the potential of such analyzers, they have yet to achieve widespread adoption for liquid chromatography/mass spectrometry (LC/MS) applications as they have lacked an effective way of interfacing to continuous ion sources.

This work aspires to resolve the limitations of previous approaches by developing a novel Astral™ mass analyzer. It is implemented in a hybrid mass spectrometer in combination with a quadrupole mass filter and the Orbitrap analyzer. The properties of the instrument are described in detail, and parallelized methods of operation are shown that employ the Orbitrap acquisition for full scans, and the Astral analyzer for MS/MS measurements to leverage the strengths of each analyzer. Experiments have been performed in accordance with standard proteomics workflows and compared against the results of current state-of-the-art instrumentation.

## Experimental Methods

### Chromatography and Sample

Instrument characterization was carried out with infused Pierce™ Flexmix™ calibration solution, which includes caffeine, the MRFA peptide, and Ultramark. LC/MS applications were performed with Pierce Hela digest ranging in amount from 250pg to 2μg, and a 3-proteome mixture of human, E.Coli and yeast digests mixed at different ratios.

Liquid chromatography was performed using a Vanquish™ Neo UHPLC system, operated either in a direct injection or trap-and-elute configuration. Sample was injected via autosampler and separated on either an Easy-Spray™ PepMap™ Neo UHPLC 150 μm x 15 cm or 50 cm low load- or 110 cm μPAC™ HPLC columns, respectively. Various gradient lengths were used to separate samples before introduction to the mass spectrometer. The instrument was operated in Data Independent Acquisition (DIA) mode and acquired raw data files were processed with Proteome Discoverer™ software using the CHIMERYS™ intelligent search algorithm^12^, or Biognosys Spectronaut™^17^. In low load experiments, the FAIMS Pro Duo™ interface was used to reduce singly charged background ions admitted to the mass spectrometer^15^.

### Mass Spectrometry

Experiments were performed using a prototype Orbitrap Astral mass spectrometer, a schematic representation of which is shown in Figure 1. Electrosprayed ions are admitted into the first ∼4 mbar vacuum region through a heated steel transfer tube of rectangular cross-section, captured by an ion funnel, and passed via quadrupole ion guides through a series of differentially pumped regions to the hyperbolic-rod quadrupole mass filter.

**Figure 1.**
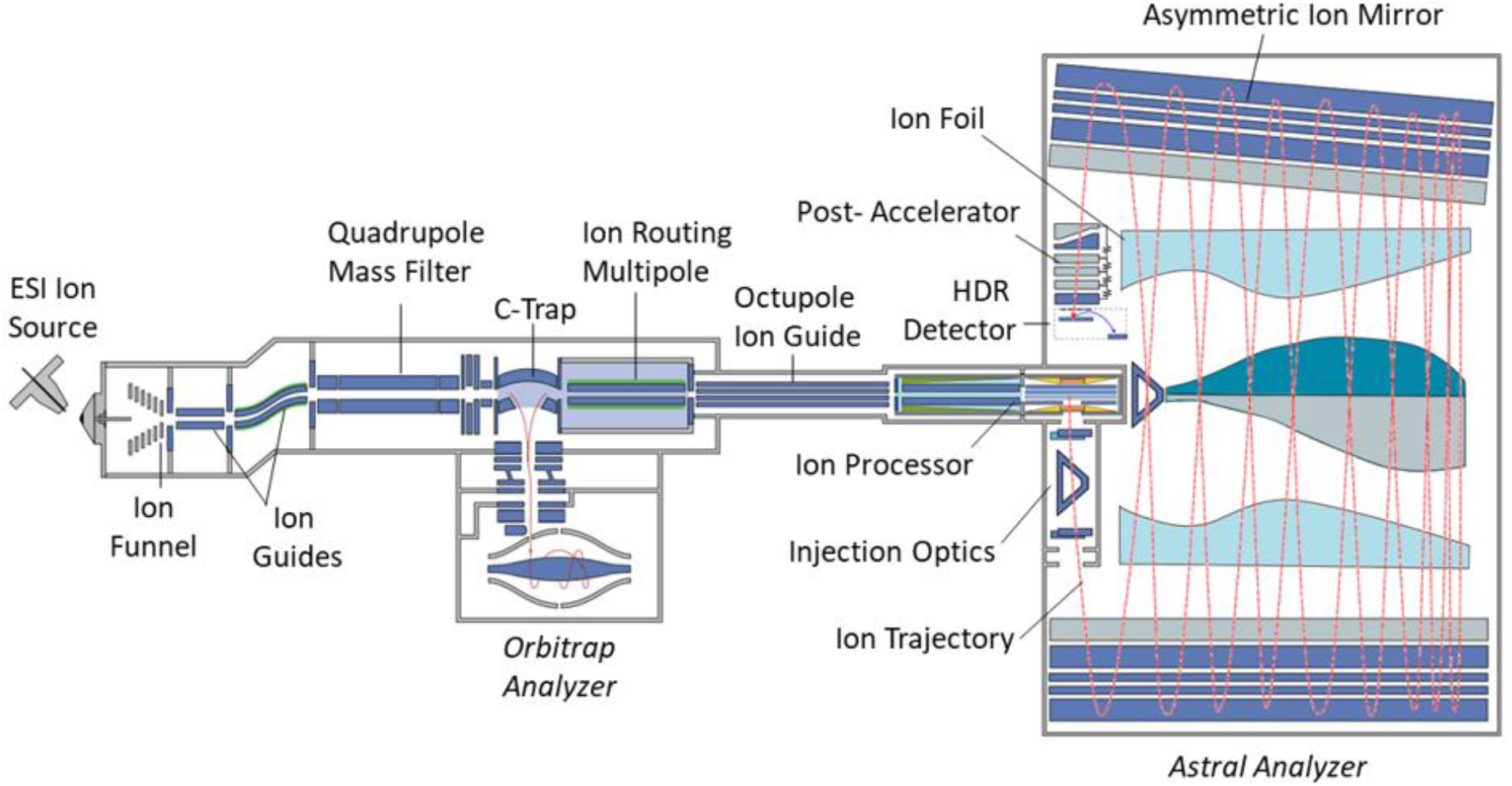
Schematic representation of the mass spectrometer.

For a full mass scan, a wide range of ions pass through the mass filter into the C-Trap, with the admission controlled by a gate between them. In normal operation, the ions are not stored in the C-trap but pass further to the Ion Routing Multipole (IRM) for trapping within a gas-filled PCB-mounted quadrupole ion guide. Inside the IRM, the excess kinetic energy of the ions is quenched in the buffer gas (ca. 10^−2^ mbar nitrogen). A series of electrodes apply a gradient voltage along the IRM to drive ions back to the C-Trap through the buffer gas^28^, where the ion packet is further collisionally cooled, compressed, and ejected into the Orbitrap analyzer via a system of lenses and deflectors. The high-field Orbitrap analyzer^29^ operates at approximately 2 Hz repetition rate with the 4 kV central electrode voltage, enabling 240K resolving power at *m/z* 200 using the Enhanced Fourier Transform (eFT™) signal processing method^30, 31^. This operation can be modified to rates as fast as 40 Hz with a resolving power of 7,500 at *m/z* 200 or 1 Hz with a resolving power of 480,000 at *m/z* 200.

For Astral MS/MS spectra, a precursor isolation window is set by the quadrupole mass filter, either identified from the full mass scan (Data Dependent Acquisition, DDA) or pre-defined by the acquisition method (Data Independent Acquisition, DIA). Ions of an isolated *m/z* range pass through the C-Trap to IRM. Contrary to the full mass scan mode, the IRM axial gradient is reversed to drive ions to the distant end, where they are blocked by a trapping voltage on an exit aperture. Therefore, the ions are trapped and stored at the far end of the IRM during a specified accumulation period, after which the positive trapping voltage is reduced to a negative transmitting level and the ions are released into a 350 mm-long octapole ion guide with ∼5.5 eV kinetic energy per unit charge.

The octapole transmits ions to the Ion Processor, represented by a dual pressure region linear quadrupole ion trap operated at 3.8 MHz RF and having the inscribed radius r_0_= 2mm. The first section of the Ion Processor is a high-pressure region fed with nitrogen by a peek capillary and operates at 10^−2^ mbar. Here the ions are accelerated to a higher energy and undergo higher-energy collisional dissociation (HCD)^32^. Wedge-shaped DC electrodes mounted between the RF electrodes create a DC gradient that drives the fragment ions to the far end of the high-pressure section where they accumulate and thermalize. A rise of the DC offset of the high-pressure section pushes the ions to the following low-pressure section. The two sections are driven with phase-locked RF supplies, which minimizes harmful fringe fields at the interface and allows ion transfer at a moderate kinetic energy ca. 1 eV per unit charge. The low-pressure section is separated by a gas conductance restriction which allows its operation with a pressure of 2-4×10^−3^ mbar, a good compromise between thermalization time and avoidance of unwanted collisions during orthogonal extraction to the Astral analyzer.

The low-pressure section of the Ion Processor contains one pair of RF rods that are split longitudinally, creating an equatorial space within which auxiliary DC electrodes are mounted, which defines an axial potential well to store and thermalize the ions in the middle of the low-pressure section in front of a slot for subsequent orthogonal ejection^33^. This accumulation process allows for very high utilization of the ion beam, compared to the poor duty cycle and transmission losses typical of orthogonal accelerators. The offset of the entire low-pressure region is lifted to 4 kV, the RF waveform is rapidly quenched, and a 500 V/mm DC pulse is applied across the trap to extract the ions through the slot into the Astral analyzer. When an ion packet is ejected from the low-pressure section and starts its motion through the analyzer, the next ion packet is already being prepared in the high-pressure section to ensure maximal instrument utilization.

The extracted ion packet is accelerated to 4 kV as it passes a grounded aperture before being shaped by a pair of lenses and an electrostatic prism comprising the Injection Optics. The tabletop-sized Astral analyzer is of a form previously described by Grinfeld and Makarov^26^. It features a pair of elongated Asymmetric Ion Mirrors, between which ions perform multiple oscillations while slowly drifting down the mirrors’ length due to the initial inclination of trajectories^24-25,34-36^. After the first reflection, the inclination angle of ion packets is adjusted by the second electrostatic prism to the optimal value of about two degrees.

The Asymmetric Ion Mirrors are designed to be slightly converging towards each other, making the ion drift decelerate over the course of the first 12-13 oscillations towards the distant end of the mirrors. The drift is eventually reversed by a returning electrostatic potential, and over the following 12-13 oscillations the ions drift back to the second electrostatic prism of the injection optics. While making a complete set of 24-26 oscillations, the ion packets are separated according to their mass-to-charge ratios and, being re-focused spatially, arrive at a High Dynamic Range detector located at the proximal end. The “Ion Foil” electrodes are mounted between the mirrors, above and below the ion path, and are biased with a small tunable potential between 0 and -20 V. The precise shapes of the Ion Foil electrodes serve to compensate for the temporal aberration induced by the mirror asymmetry and improve the quality of the spatial focus at the detector, as well as to compensate for mechanical misalignment of the mirrors. This makes ion motion isochronous both along and across the mirror system.

The combination of gridless design with spatial focusing enables a high transmission of ions throughout the analyzer. The asymmetric geometry of the analyzer as well as its high transmission are reflected in the Astral abbreviation which stands for ASymmetric TRAck Lossless analyzer. To reduce ion losses to collisions with the residual gas on the long track, the main chamber of the Astral analyzer is maintained at a pressure below 10^−8^ mbar by a 6-port split-flow turbopump (Pfeiffer, Asslar, Germany).

The High Dynamic Range (HDR) detector (El-Mul Technologies Ltd) is a novel device incorporating a conversion dynode, a magnet for focusing of secondary electrons, a scintillator, light guide, and a photomultiplier tube (PMT). Thanks to optical coupling, the PMT is hermetically sealed, which excludes contamination of dynodes by residual gases. The detector’s compact design reduces noise and space-charge effects while ensuring high sensitivity and dynamic range with a long detector lifetime.

A post-accelerator, implemented as a stack of slit-shaped apertures, is mounted in front of the detector to deliver an extra 10 kV acceleration voltage to the ions before hitting the conversion dynode; the total impinge energy thus being as high as 14 keV per charge. This enhances the secondary electron yield for more efficient detection of ions, particularly at high m/z, maximizing probability of successful conversion. The secondary electrons are further accelerated to 6 keV before being converted to photons in the scintillator, and then converted back to electrons and amplified in the PMT. The detector signal is split into a two-channel preamplifier with a 10x difference in gain. to further enhance dynamic range.

In total, the ions undergo 24-26 reflections at each mirror, with a total asymmetric track length of over 30 meters, and an arrival time around 800μs for m/z 524 MRFA peptide ions.

Repetition rate is limited by numerous factors, but most important are the accumulation time (fill time) for sufficient ions to be accumulated for analysis, the time taken by power supplies to dynamically adjust during scan execution, and the time required for ions to traverse the mass spectrometer. A complex instrument with multiple sequential stages can easily bring the total operation cycle well above 10ms even before consideration of ion accumulation time. For this reason, each stage here has been highly parallelized with multiple ion packets simultaneously being handled to bring the maximum repetition rate to above 200Hz.

Figure 2 shows a representation of the operational sequence for an Astral analyzer MS/MS scan, with the time required for each stage. The time for the ion source, including the interface ion guides, is given as variable depending on the difference in *m/z* between the targeted ion packet and its predecessor. For the small mass steps common in DIA acquisition there is little difference in transmission before and after voltage adjustment. Larger mass jumps, as in DDA experiments, incur an extra penalty for ions to travel from the capillary to the quadrupole. The accumulation of ions in the IRM, rather than directly to the Ion Processor, is added to maximize allowed ion accumulation/fill time through parallelization with the Ion Processor’s operation. No non-parallelizable series of stages exceeds >4.5ms in DIA, allowing 200Hz repetition rate.

**Figure 2.**
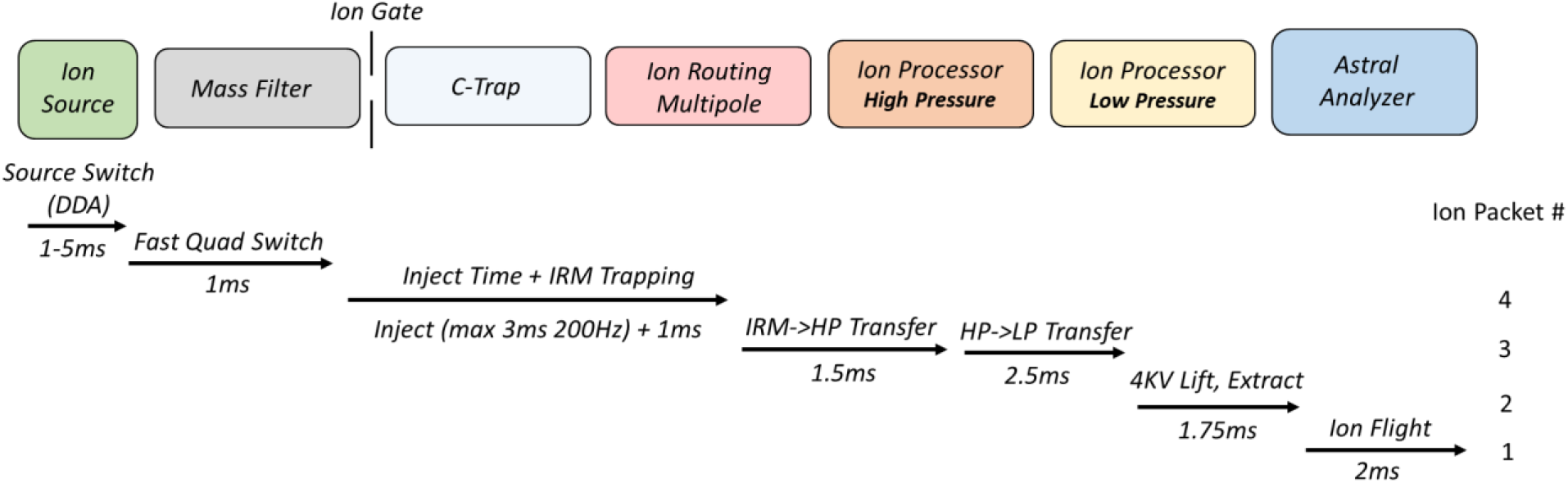
Timing sequence for 200Hz parallelized ion transfer of ions from source to Astral analyzer, with trapping points in IRM, and both high and low-pressure extraction trap regions.

### LC-MS Analysis Method

The hybrid acquisition cycle is drawn schematically in Figure 3. For both DIA and DDA acquisition, a full mass range of ions was first injected into the Orbitrap analyzer, and then a series of MS/MS scans performed whilst the Orbitrap analyzer acquired the next full scan. Full mass scans were also performed in the Astral analyzer at regular intervals for automatic gain control (AGC^37^) (so called “fluxscans”) to measure ion current and then set appropriate accumulation times for the Orbitrap full mass scans, precursor peak masses and intensities from which were then used to control isolation targets and accumulation times for MS/MS. The AGC target level was set to 100% for full mass scans, equivalent to very approximately 50,000 charges for full mass scans. For MS/MS 100% AGC relates to around 10,000 charges, however with a maximum inject time of 3ms, higher MS/MS AGC settings did not substantially slow the system down and thus had relatively little drawback. Accumulation time for MS/MS scans varied from 0.03-3ms, but mostly reached this limit. Orbitrap resolution was set to 240K, equivalent to 2 Hz full-MS acquisition.

**Figure 3.**
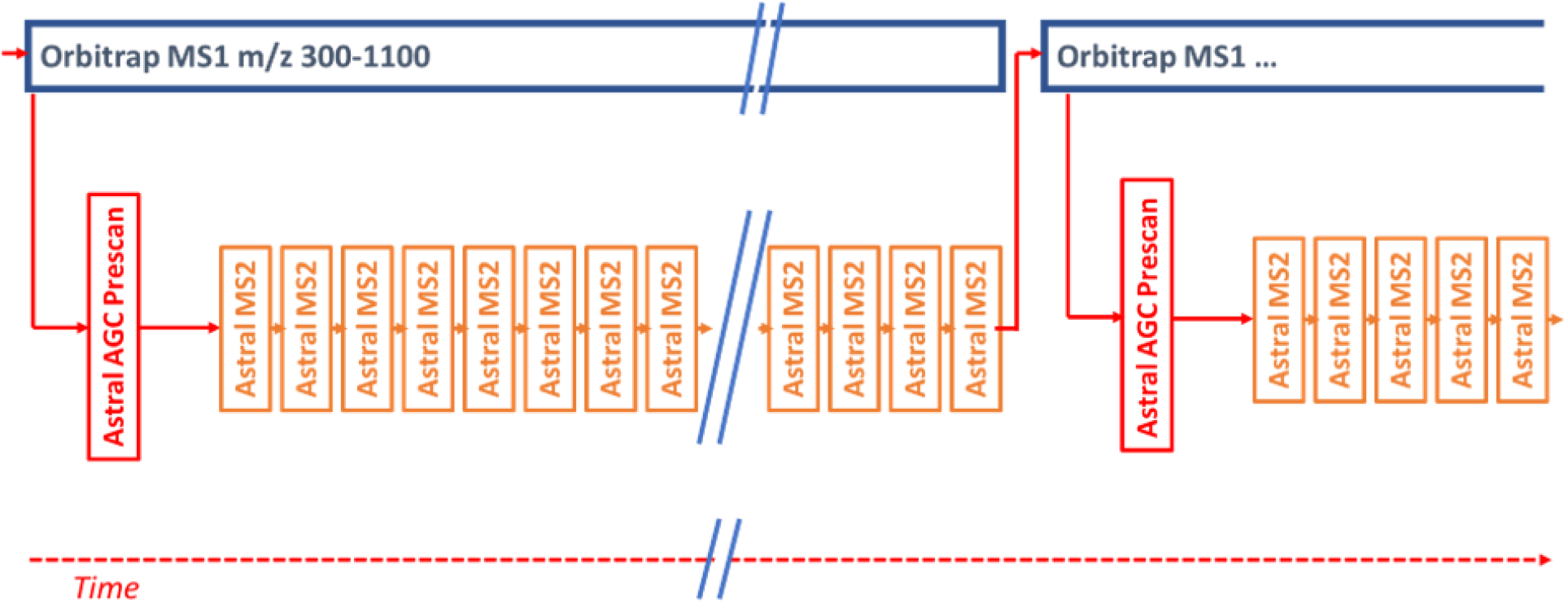
Representation of parallelized acquisition method, with Orbitrap full MS precursor detection scans overlapped with Astral MS2 identification scans.

## Result and Discussion

### Instrument Characterization

Various properties of the Astral analyzer were characterized with calibration solution. Figure 4 shows the resolution and mass deviation of averaged Flexmix peaks across a 190-3000 m/z range, with 5000 and 50,000 total ions detected per shot. Number of ions was determined by measuring the signal response of single ions and dividing the sum of the detected peak intensities accordingly. Increased ion loads generate space charge effects within the Ion Processor, expanding the trapped ion cloud, whilst also causing additional expansion of extracted ion packets as they traverse the analyzer. Unsurprisingly average resolution was reduced from above 100K to ∼90K at the higher ion load. More severe is the loss of resolution for individual very intense analyte species. The mass shift imposed at high ion load was also less than 1ppm for all ions, with no applied correction to mass position for peak intensity.

**Figure 4.**
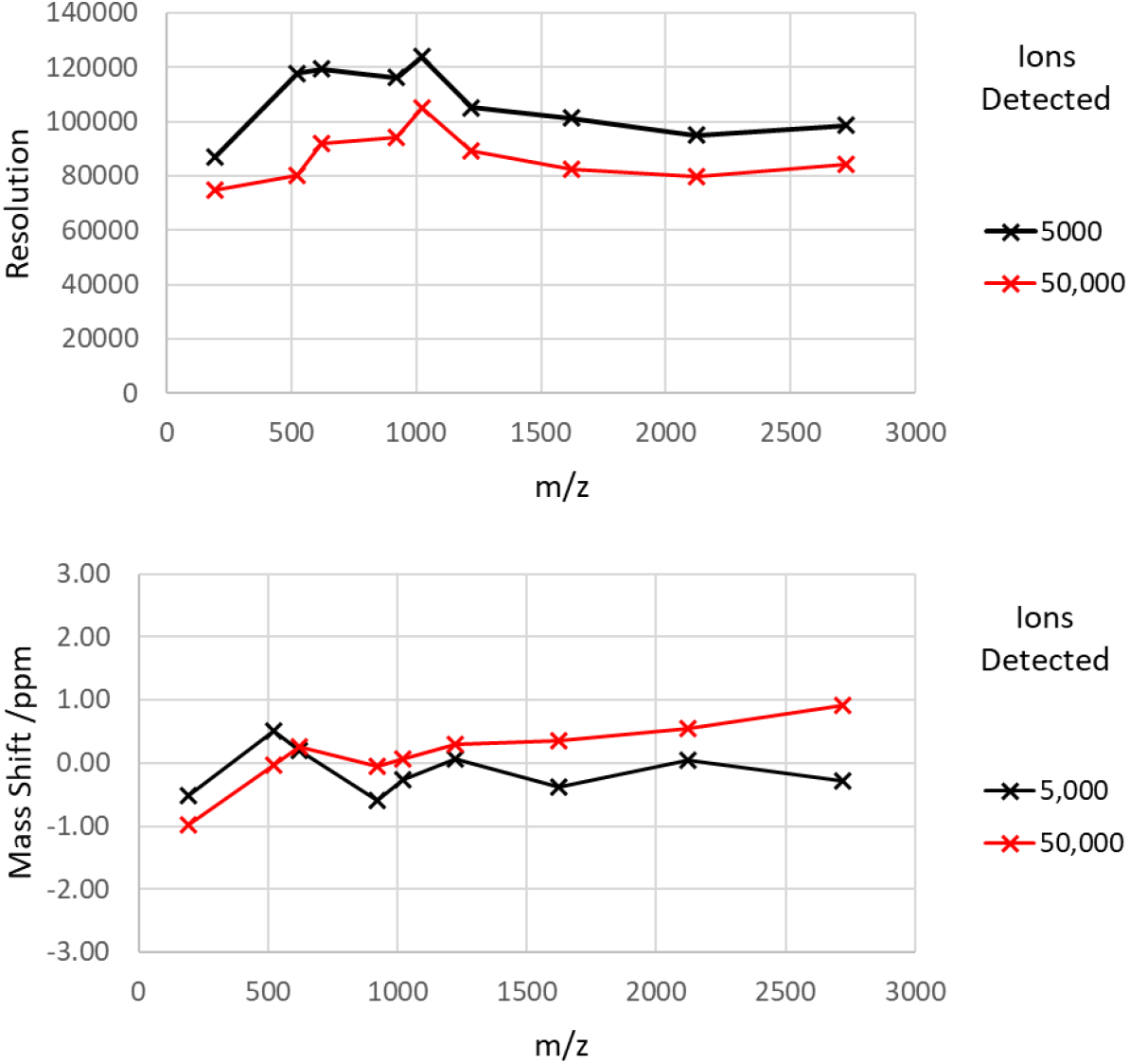
Resolution and m/z measurement shift of calibration mixture ions at 5000 and 50,000 ions detected per shot.

For rapid spectra acquisition, Astral methods follow the same strategy as the Orbitrap acquisition, i.e. that every scan forms a spectrum, taking advantage of high transmission and signal-to-noise ration to avoid time-consuming averaging. However, in this case mass accuracy is determined by shot-to-shot jitter. This was measured by gathering single shot peak data over 60 seconds and calculating the standard deviation σ. The 3σ jitter is shown in Figure 5 and was largely observed to be <1.5ppm (and thus 1σ generally <0.5ppm), with the exception of m/z 1022 and 1922, which were comparatively low intensity peaks with high statistical error.

**Figure 5.**
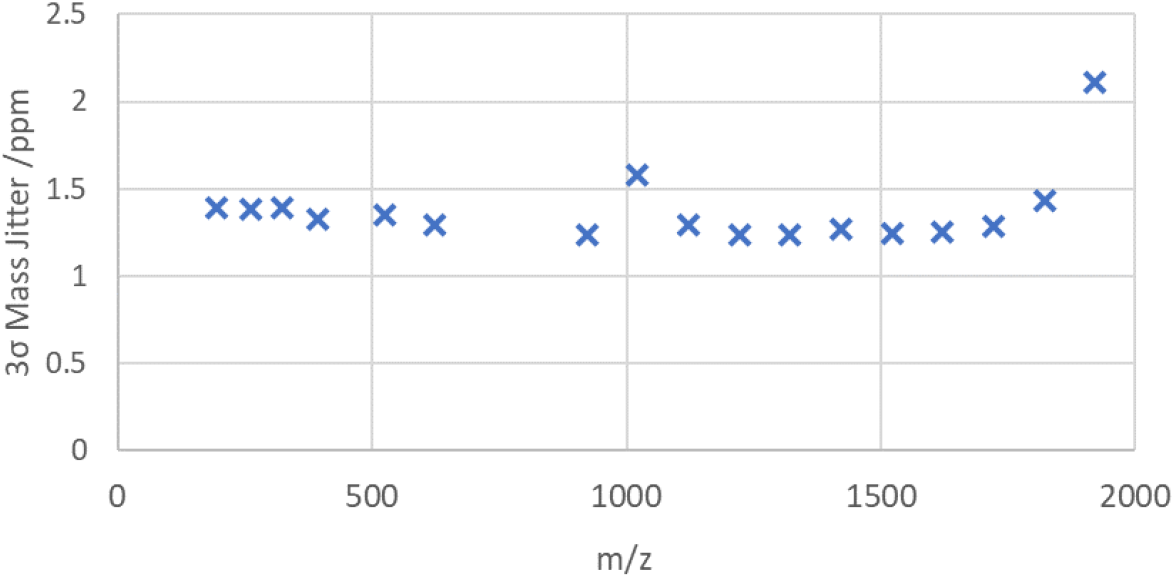
3-sigma level mass jitter for single shot repetitions.

To control mass measurement error caused by drift of the analyzer mechanics or applied voltages, an intermittent internal calibration method was developed, whereby an internal calibrant ion (in this case the +1 ion of fluoranthene at m/z 202) was injected regularly and the measured time added to a moving average value used to proportionally correct to arrival times of all analyte peaks. An injection once every 10 seconds with a 10-point moving average was found to be sufficient for this purpose, though even less frequent injection such as every 5 minutes may also be used. Whilst this has no benefit for control of jitter, as with conventional internal calibration applied to every mass spectrum, it instead has the advantage of having a negligible impact on overall repetition rate.

Figure 6 shows the application of this method to the recording of Flexmix Ultramark 922 peaks over a 70-hour period, where the corrected measurement remains largely within 0.5ppm (0.13ppm standard deviation). 50x averaging was used to suppress shot-to-shot jitter. Combining this measurement with the estimation for shot-to-shot jitter, and some allowance for quality of calibration, implies that the vast majority of peaks should be measured to well within 3ppm mass accuracy. The uncorrected drift, without even temperature compensation, is driven over a 2.5ppm range largely by a 0.8K laboratory thermal drift. With temperature sensors mounted in the vacuum chamber and the grounded mirror electrode, it is also possible to compensate for thermal expansion of the analyzer by measuring the temperature change and applying a suitable constant (3.5ppm/K) and delay function. Figure 7 shows a second similar experiment, separately recorded, showing the influence of a 3ppm/K temperature compensation applied with a 30-minute weighted average delay. Here a 2.5ppm drift, driven by a 1K temperature shift, was reasonably well corrected for.

**Figure 6.**
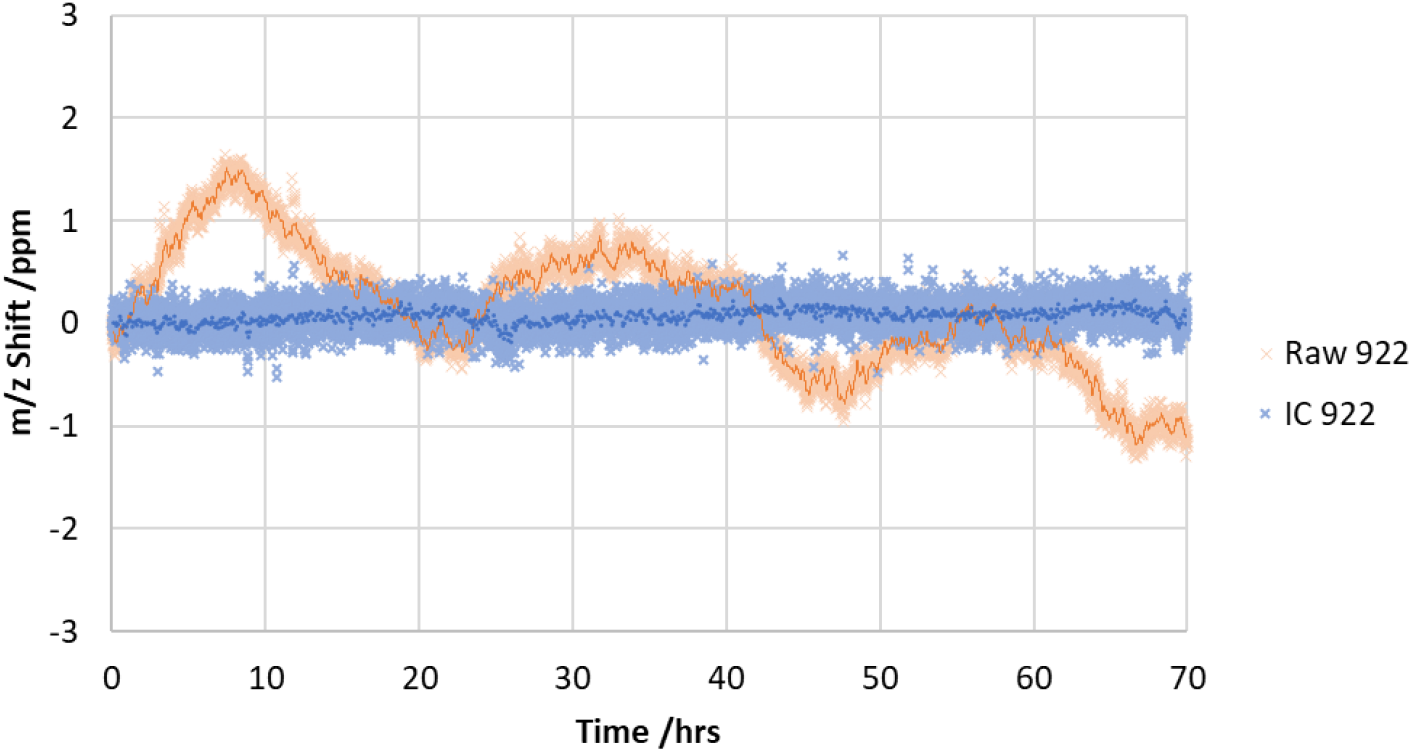
Mass drift of Ultramark m/z 922 over 70 hours, as raw unmodified signal and with an intermittent internal calibrant correction.

**Figure 7.**
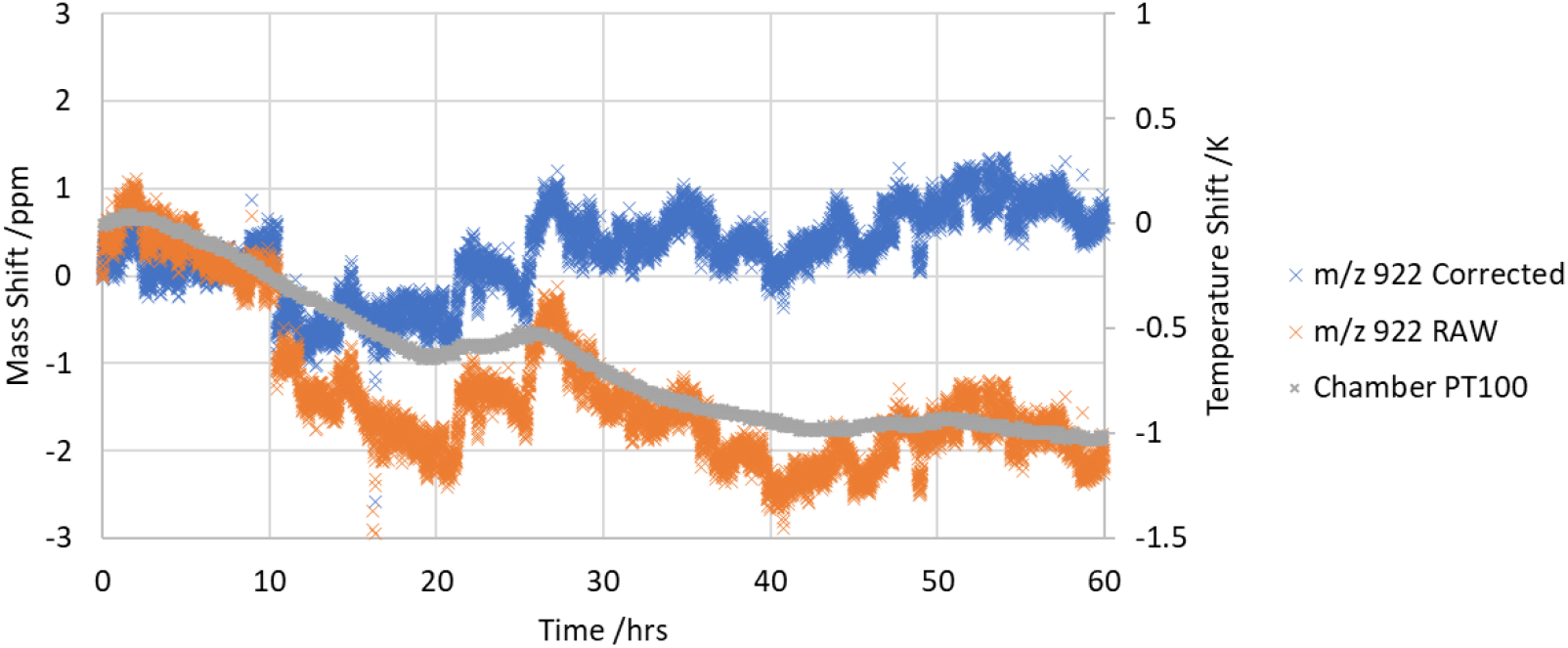
Mass drift of Ultramark m/z 922 over 60 hours, as raw unmodified signal and with an applied 3ppm/K temperature compensation. Temperature drift at the chamber corner is also shown.

As described, an important consideration for fast LC-MS experiments is the Astral analyzer’s repetition rate and duty cycle. This latter property may be defined as the proportion of the scan interval over which that the ion trap may spend accumulating ions from the source, prior to pulsed extraction into the analyzer. A scan of ion accumulation time was performed, and the repetition rate monitored and drawn graphically in Figure 8. Below 2.5ms the accumulation time has no impact on the repetition rate, but above this repetition rate falls precipitously as the accumulation time comes to dominate the operational cycle. At 200Hz the maximum accumulation time was just over 3ms, equating to a 60% duty cycle, though it may in future be possible to increase this via pre-accumulation of ions in the ion guide ahead of the quadrupole, as previously shown for the Orbitrap Exploris mass spectrometer^17^. In a second experiment, the scan rate was recorded whilst the instrument was run in a DIA mode of operation, with a fixed 3ms accumulation time. Here the scan rate was recorded consistently above 200Hz, as also shown in Figure 8. This is all very suitable for DIA experiments with small m/z transitions, however precludes extra overheads for large transitions (DDA), that require an additional 2ms to adjust the ion source.

**Figure 8.**
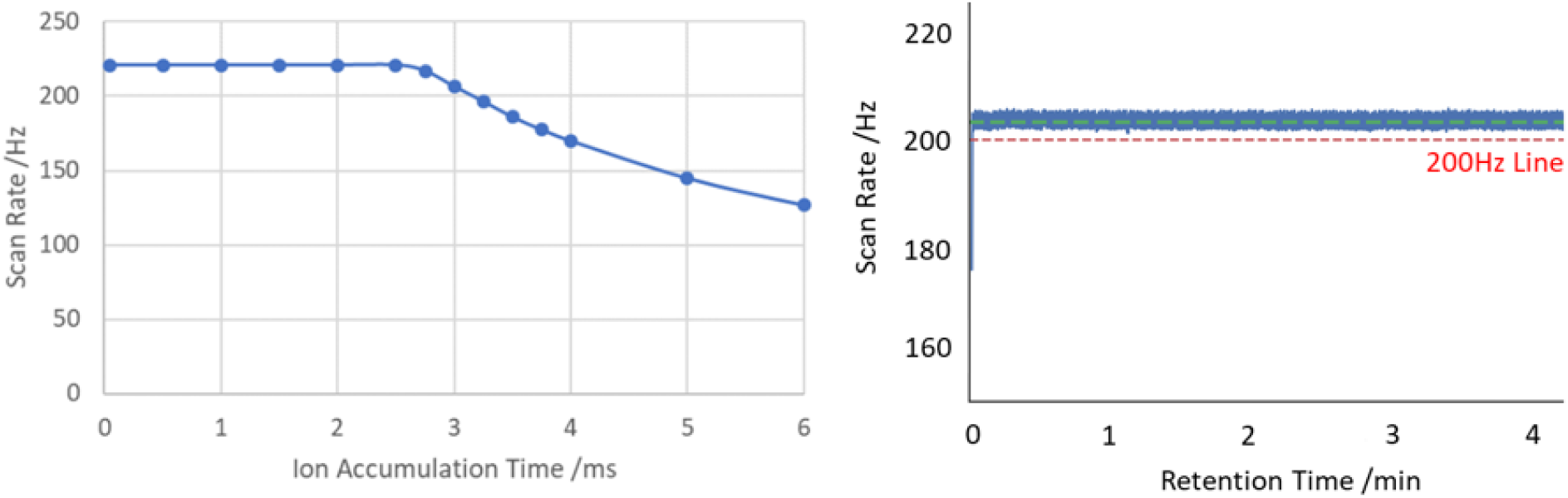
Left: Astral repetition rate with varying ion trap accumulation times, where acquisition drops below 200Hz with >3ms accumulation time. Right: Scan rate recorded whilst running DIA method with fixed 3ms ion accumulation time.

The sensitivity of the Astral analyzer was compared to that of the Orbitrap analyzer by comparison of ion number counts within sequential injections to each analyzer, of differing m/z Flexmix ions. Orbitrap ion numbers were estimated from signal to noise (S/N) measurement of multiple ion peaks, compared to earlier S/N measurement of multiply charged single ions^38^. Highly charged ions, such as intact proteins, produce intense single ion peaks that the Orbitrap analyzer is capable of detecting^39^, allowing estimation of the number of charges at the noise threshold; approximately 2.6 for an Orbitrap Exploris MS running a 256ms transient (resolution 120K) with eFT signal processing.

Astral analyzer peak intensities were also converted to ion numbers by comparison with measurement of single ions of MRFA at m/z 524. A mass dependent correction was also applied based on additional single ion measurements of the other Flexmix m/z ions, as the conversion of ions to secondary electrons at the detector is strongly m/z dependent^40^. The results, shown in Figure 9, even without correction indicate substantially greater transmission of ions into the Astral analyzer, perhaps double at higher m/z, and consequently higher sensitivity. At the lowest m/z however the trend is reversed; the reason for this is thought to arise from ion transport losses between IRM and Ion Processor, as fragmentation spectra did not produce such discrimination. It is recognized that the smaller inscribed radius of the Ion Processor vs the C-Trap results in more difficult trapping conditions at low m/z. It is also likely that the assumption of constant Orbitrap noise level does not hold over this wide mass range.

**Figure 9.**
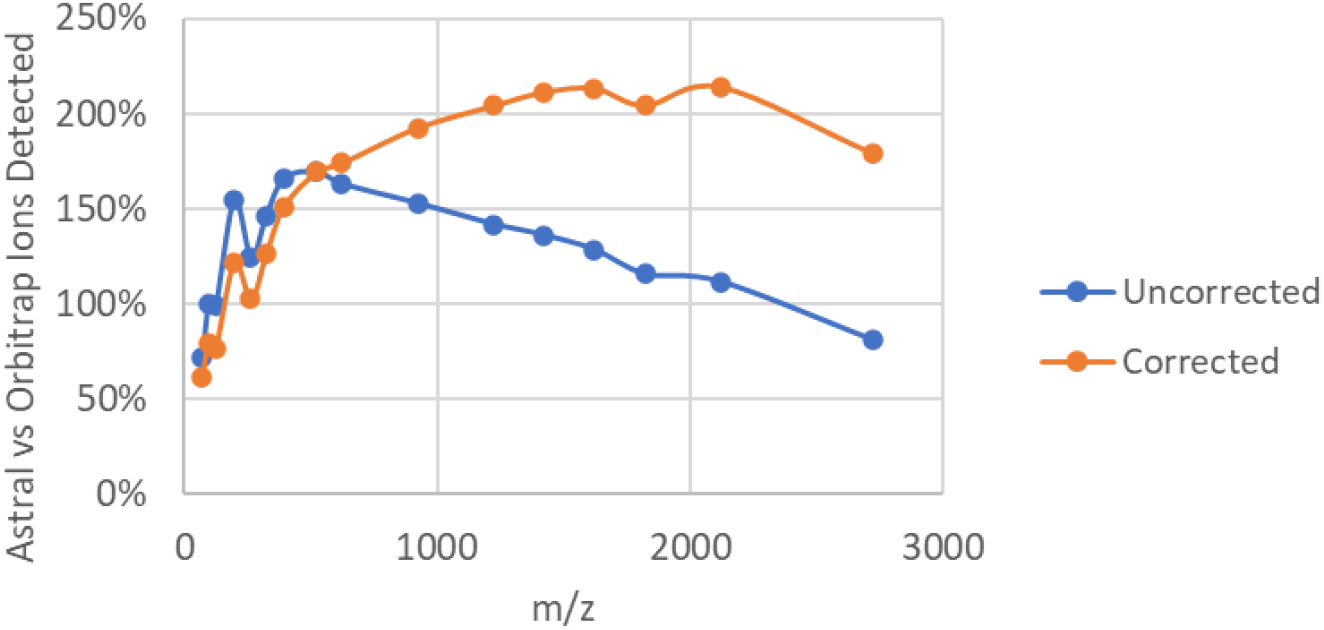
Relative number of ions detected in Astral vs Orbitrap analyzers.

Figure 10 reproduces a typical peptide MS/MS spectrum, in this case infused MRFA, in the Astral analyzer, along with a typical MRFA raw peak profile recorded by both detector channels, acquired separately. The fragment mass positions in the Astral spectrum matched that of an equivalent Orbitrap scan, but notably the peak resolution was ∼100,000 except at m/z below ∼150, where resolution remained high but began to clearly drop off. An Orbitrap DDA or DIA method would typically run at resolution values of 7.5 or 15K at m/z 200. A significant advantage of the Astral analyzer over the Orbitrap analyzer is the detection of single ions, rather than requiring >10 ions for a detectable peak in MS/MS acquisition. Figure 10 also shows a side-by-side comparison of a single-shot MRFA fragment spectrum in both analyzers, with the ion accumulation time targeting ∼1000 ions detected in the Astral analyzer. The difference in the quality of the spectrum is very clear, with an order of magnitude difference in signal-to-noise, and 132 peaks detected vs only 30 in the Orbitrap mass spectrum. For identification of low-level analytes this is expected to be highly advantageous, and essential when cycling through precursors rapidly with limited ion accumulation time.

**Figure 10.**
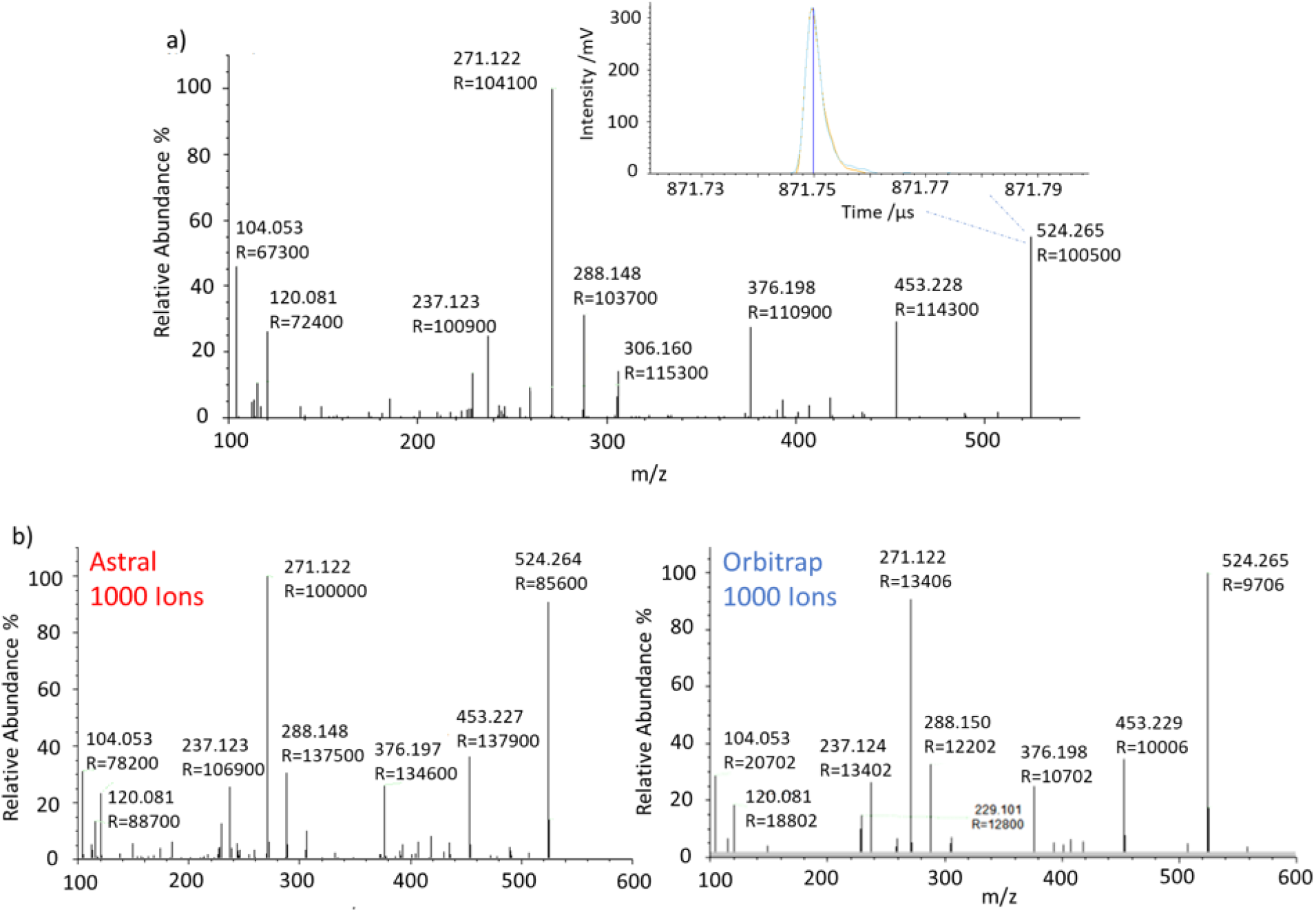
Top) MS/MS spectrum of MRFA Peptide in Astral analyzer. Bottom) Side by side comparison of MRFA MS/MS spectra in Astral and Orbitrap analyzers, with only 1000 ions detected.

### Assessment of Proteomics Application Performance

#### DIA Throughput and Depth of Analysis

In a first experiment, 200ng Pierce HeLa digest was injected and separated on an Easy-Spray PepMap, 150 μm x 15 cm column with 300, 180, 100, and 60 samples per day (SPD) methods (3.6, 5.5, 11, and 20.1-minute gradients corresponding to 4.8, 8, 14.4, and 24minute experimental cycles respectively). A second experiment used 1000ng load over slower 30 SPD runs, with separation on a μPAC Neo 110 cm column. The AGC target was set to 500%, ∼50,000 charges restricted by a maximum inject time of 3ms for the first experiment and 2.5ms for the second, the full-MS mass range to 380-980, and the MS/MS range to 150-2000 with 2 Th windows. Figure 11 shows the results obtained with Chimerys, reporting protein and peptide identifications filtered to <1% peptide FDR and <=1% protein FDR. Three technical replicates for each SPD method were processed together using Chimerys with match-between-runs.

**Figure 11.**
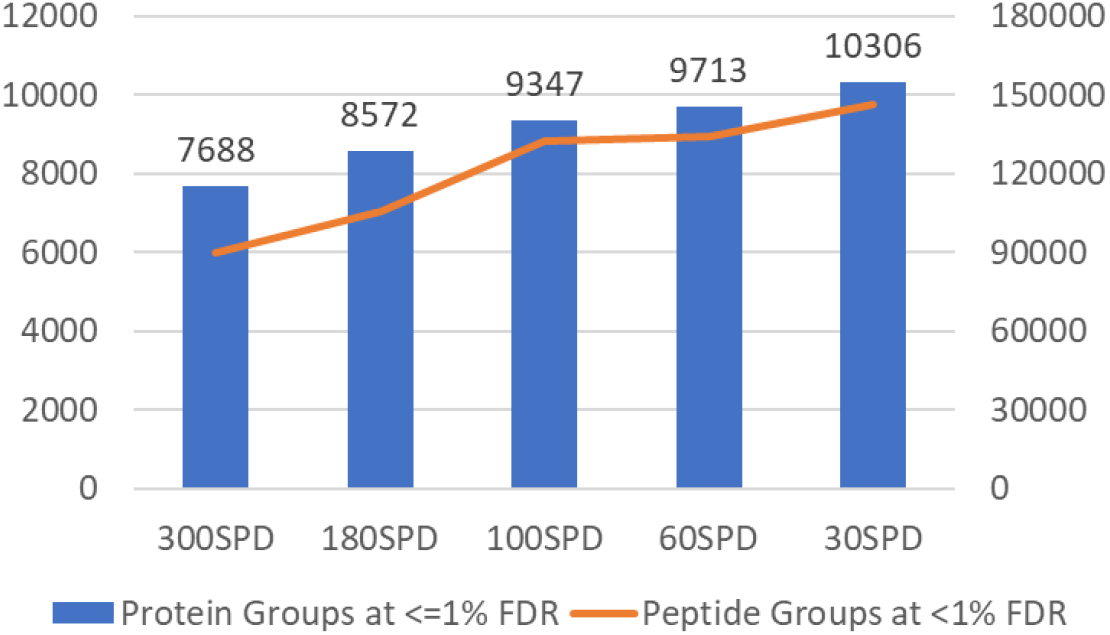
Single shot HeLa DIA protein groups and peptide groups IDs. Variation in experiment length from 300-30 samples per day. 200ng HeLa digest was injected for 300, 180, 100 and 60 SPD runs and 1000ng HeLa digest for the 30 SPD runs.

The depth of analysis is extremely high for a single shot method, evidenced by the 10,306 protein groups identified over a 48 min run. As might be expected, there remains a balance between throughput and analytical depth, but even the 4.8-minute experiment shows >7,600 protein groups identified, rising to >9,000 for the same quantity analyzed over 14.4-minutes. A rough comparison is that Orbitrap-only methods typically require at least 4x the LC separation time to produce similar depth of analysis to the hybrid Orbitrap/Astral methods.

#### Label-Free Quantitation

Quantitative performance was assessed by analysis of two 3-proteome mixed sample containing E. coli, HeLa and yeast digests at 2:3:1 (sample A) and 1:3:2 (sample B) ratio. 500ng loads were injected and separated on a 50cm μPAC Neo column over a 24-minute experiment. The mass spectrometer itself was operated in the same manner as the throughput experiments above, but for a 3ms Astral max inject time, and the output data was analyzed in Spectronaut 17 using the DirectDIA workflow.

Figure 12a shows the overall protein IDs and the proportion within 20 and 10% coefficients of variation CV. Of more than 13,000 identified proteins, the median CV was <4.7%, whilst over 72% had a CV <10% and 87% a CV <20%. Figure 12b shows average log2 ratios of intensities of the identified human, yeast, and E.coli proteins between the two 3-proteome mixtures, A and B. Theoretical A/B ratios were 1 for human, 0.5 for yeast, and 2 for E.coli proteomes. Experimentally obtained average ratios were 1.016 for human, 0.490 for yeast, and 1.993 for E.coli. Run-to-run reproducibility of quantitation was better than 1%. High quantitative performance was maintained throughout various loads and gradient lengths and remained unchanged with minor MS acquisition method variations. According to Spectronaut data, such quantitation accuracy was obtained with a median 3-4 data points per peak (FW defined as 1.7* FWHM), which corresponds to 4-6 baseline-to-baseline datapoints.

**Figure 12.**
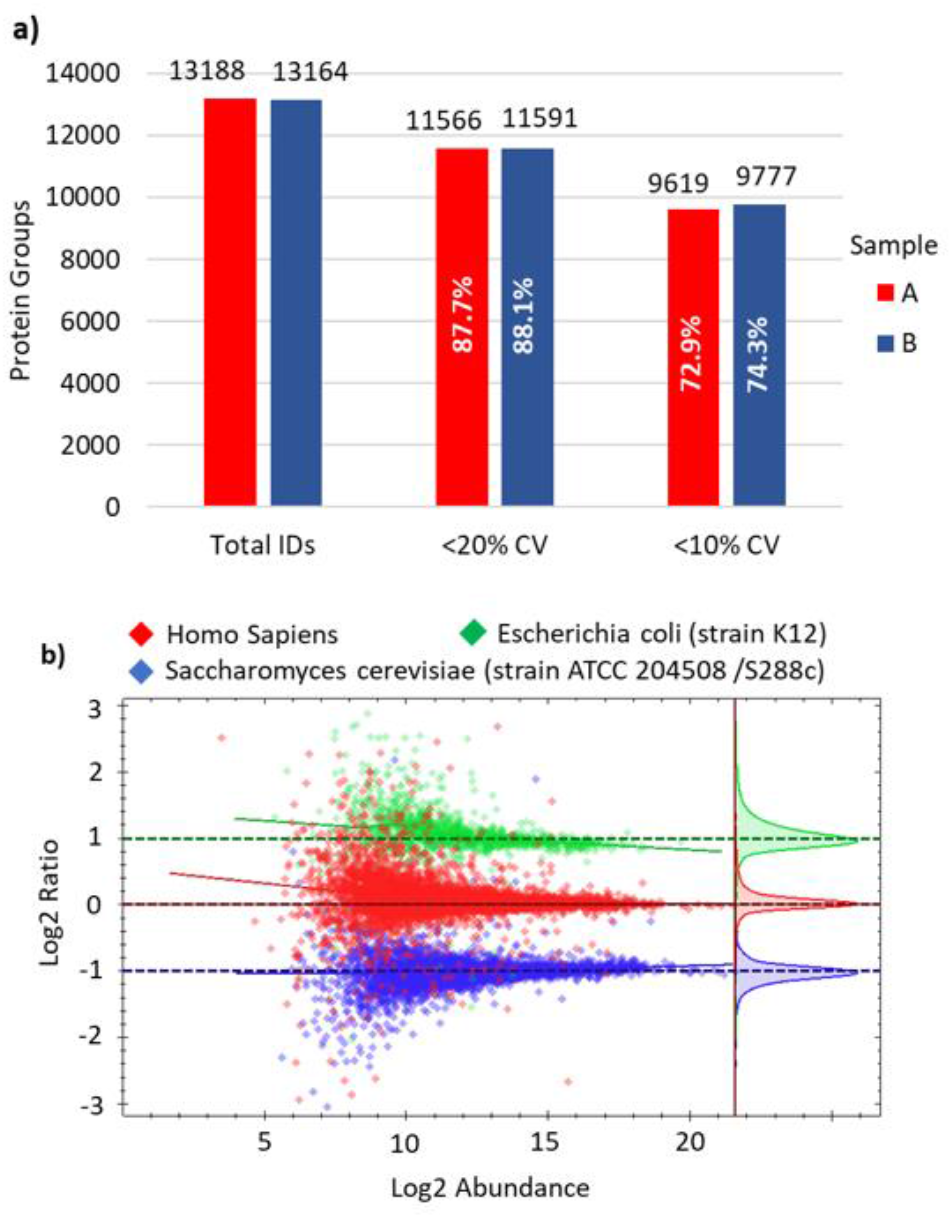
Comparative analysis of 500ng 3-proteome mixtures (E. coli, HeLa & Yeast) at 2:3:1 and 1:3:2 ratio for samples A & B respectively. a) Protein identifications at various coefficients of variation between triplicate runs. b) Average Log2 ratio of yeast and E. coli protein intensities between samples A and B.

Increasing the DIA window from 2 Th to 3 and 4 Th raised the number of Spectronaut datapoints to 5 (7-8 baseline-to-baseline) and 6 (9 baseline-to-baseline) per peak correspondingly, yet this did not influence the average quantitation accuracy. Maintaining good quantitative performance over a higher ID range is almost surprising, as greater sensitivity to smaller numbers of ions might be expected to result in greater statistical fluctuations of trace species. However, this seems not to be the case, and the mass spectrometer remains sensitive to small changes in protein amounts within complex mixtures.

#### Low loads

With the explosion of the single cell analysis field, high throughput measurement of very low amounts of protein sample have become increasingly important. Small amounts of HeLa from 0.25 to 10ng were separated over an 18-minute, 80 SPD method on a 50cm μPAC Neo low load column and measured by DIA with a full-MS mass range of 400-800. The isolation width and maximum injection time were decreased from 20Th and 40ms for 0.25ng load to 5Th and 10ms for 10ng load. The FAIMS Pro Duo interface was used with a compensation voltage of -50V to remove singly charged background ions, which prevented the ion processor becoming flooded by unwanted ions. Data was processed using Spectronaut 17, both with the library-free DirectDIA workflow and by using a spectral library generated from the series of runs.

Figure 13a shows the protein and peptide identifications over the series of runs. Even at 0.25ng, 3900 protein groups may be identified with DirectDIA, increasing to 5700 with the aid of the generated library. The longer 40ms accumulation times,, are required for such low ion loads, and the instrument cannot operate at high repetition rate. The depth of identification may then be assumed to arise from the combination of high ion transmission and single ion sensitivity. Coefficients of variation are still relatively low, with 1 ng runs producing a median CV of 8.1% with MS1 quantitation, and 8.7% with MS/MS quantitation. 87% of protein IDs have a CV of <20%, and 70% a CV <10%.

**Figure 13.**
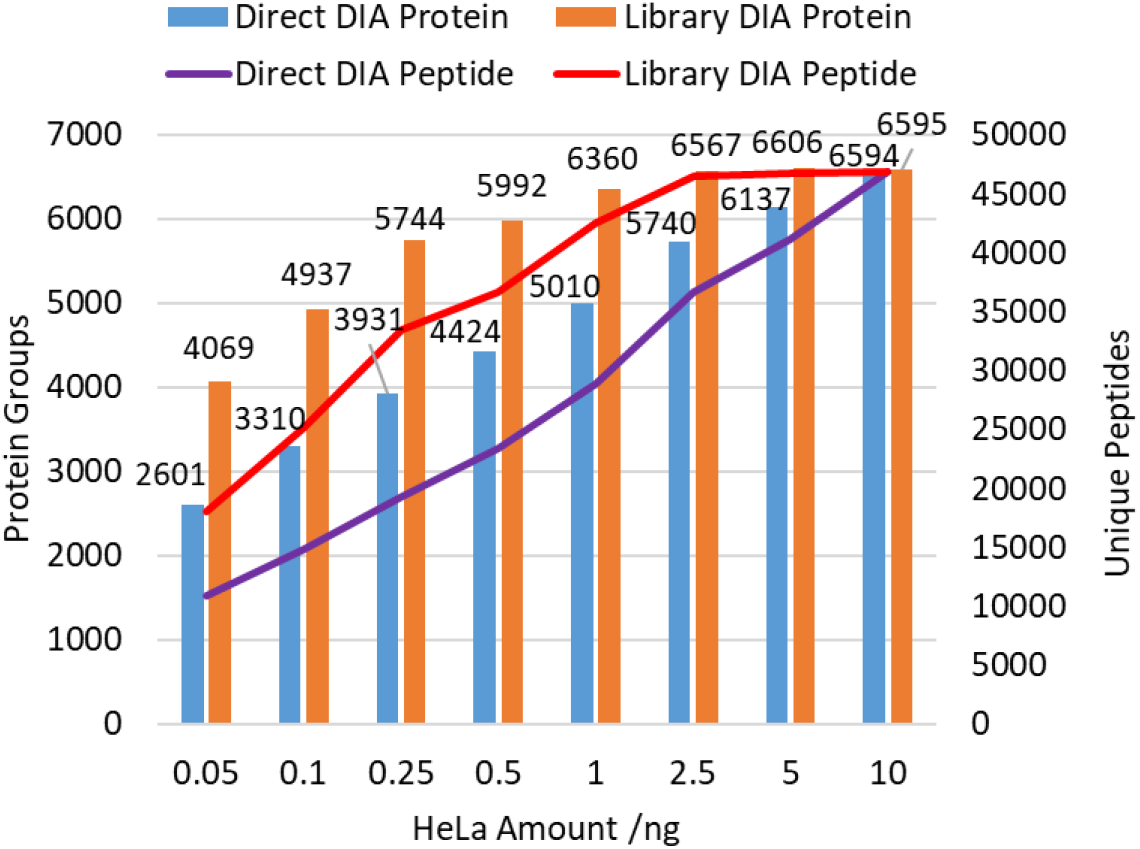
Low load HeLa 80 samples per day DIA protein and peptide identifications, obtained with and without a generated library.

## Conclusion

The Orbitrap/Astral dual analyzer mass spectrometer was characterized via LC-MS shotgun proteomics methods with a parallelized, hybrid mode of acquisition. From low to high sample loads, it was found to produce depth of analysis at higher throughputs than have hitherto been accessible. Analytical performance was thought to be enabled by the high scan rate, transmission, and single ion detection of the Astral analyzer, which allows both a high rate of acquisition to cover rapidly eluting peptides, and the sensitivity to measure low-abundance analyte species. It is anticipated that this new technology will make an important contribution the field of biological mass spectrometry.

## Acknowledgements

The authors would like to acknowledge the advice and support of colleagues throughout Thermo Fisher Scientific, without whom we would be lost, as well as academic and industrial collaborators who have been so vital to both building the next generation mass analyzers and getting the best out of them.

## References

1 Zhang, Z., Wu, S., Stenoien, D. L., & Paša-Tolić, L. (2014). High-throughput proteomics. Annual review of analytical chemistry, 7, 427–454.

2 Eng, J. K., McCormack, A. L., & Yates, J. R. (1994). An approach to correlate tandem mass spectral data of peptides with amino acid sequences in a protein database. Journal of the American society for mass spectrometry, 5(11), 976–989.

3 Perkins, D. N., Pappin, D. J., Creasy, D. M., & Cottrell, J. S. (1999). Probability-based protein identification by searching sequence databases using mass spectrometry data. ELECTROPHORESIS: An International Journal, 20(18), 3551–3567.

4 Demichev, V., Messner, C. B., Vernardis, S. I., Lilley, K. S., & Ralser, M. (2020). DIA-NN: neural networks and interference correction enable deep proteome coverage in high throughput. Nature methods, 17(1), 41–44.

5 Zolg, D. P., Gessulat, S., Paschke, C., Graber, M., Rathke-Kuhnert, M., Seefried, F., … & Frejno, M. (2021). INFERYS rescoring: Boosting peptide identifications and scoring confidence of database search results. Rapid Communications in Mass Spectrometry, e9128.

6 Michalski, A., Cox, J., & Mann, M. (2011). More than 100,000 detectable peptide species elute in single shotgun proteomics runs but the majority is inaccessible to data-dependent LC− MS/MS. Journal of proteome research, 10(4), 1785–1793.

7 Kim, M. S., Pinto, S. M., Getnet, D., Nirujogi, R. S., Manda, S. S., Chaerkady, R., … & Pandey, A. (2014). A draft map of the human proteome. Nature, 509(7502), 575–581.

8 Wilhelm, M., Schlegl, J., Hahne, H., Gholami, A. M., Lieberenz, M., Savitski, M. M., … & Kuster, B. (2014). Mass-spectrometry-based draft of the human proteome. Nature, 509(7502), 582–587.

9 Senko, M. W., Remes, P. M., Canterbury, J. D., Mathur, R., Song, Q., Eliuk, S. M., … & Zabrouskov, V. (2013). Novel parallelized quadrupole/linear ion trap/Orbitrap tribrid mass spectrometer improving proteome coverage and peptide identification rates. Analytical chemistry, 85(24), 11710–11714.

10 Eliuk, S., & Makarov, A. (2015). Evolution of orbitrap mass spectrometry instrumentation. Annu. Rev. Anal. Chem, 8(1), 61–80.

11 Scheltema, R. A., Hauschild, J. P., Lange, O., Hornburg, D., Denisov, E., Damoc, E., … & Mann, M. (2014). The Q Exactive HF, a Benchtop mass spectrometer with a pre-filter, high-performance quadrupole and an ultra-high-field Orbitrap analyzer. Molecular & Cellular Proteomics, 13(12), 3698–3708

12 Michalski, A., Damoc, E., Hauschild, J. P., Lange, O., Wieghaus, A., Makarov, A., … & Horning, S. (2011). Mass spectrometry-based proteomics using Q Exactive, a high-performance benchtop quadrupole Orbitrap mass spectrometer. Molecular & cellular proteomics, 10(9).

13 Frejno, M., Zolg, D. P., Schmidt, T., Gessulat, S., Graber, M., Seefried, F., Rathke-Kuhnert, M., … & M. Wilhelm, CHIMERYS: An AI-Driven Leap Forward in Peptide Identification, Proceedings of the 69th ASMS Conference on Mass Spectrometry and Allied Topic, Philadeplhia PA, 2021.

14 Dorfer, V., Maltsev, S., Winkler, S., & Mechtler, K. (2018). CharmeRT: boosting peptide identifications by chimeric spectra identification and retention time prediction. Journal of proteome research, 17(8), 2581–2589.

15 Bekker-Jensen, D. B., Martínez-Val, A., Steigerwald, S., Rüther, P., Fort, K. L., Arrey, T. N., … & Olsen, J. V. (2020). A compact quadrupole-orbitrap mass spectrometer with FAIMS interface improves proteome coverage in short LC gradients. Molecular & Cellular Proteomics, 19(4), 716–729.

16 Kelstrup, C. D., Jersie-Christensen, R. R., Batth, T. S., Arrey, T. N., Kuehn, A., Kellmann, M., & Olsen, J. V. (2014). Rapid and deep proteomes by faster sequencing on a benchtop quadrupole ultra-high-field Orbitrap mass spectrometer. Journal of proteome research, 13(12), 6187–6195.

17 Arrey. T., Stewart. H. & Harder. A., Ion Pre-Accumulation for High Speed Orbitrap Exploris Operation, Proceedings of the 70^th^ ASMS Conference on Mass Spectrometry and Allied Topic, Minneapolis MN, 2022.

18 Meier, F., Brunner, A. D., Koch, S., Koch, H., Lubeck, M., Krause, M., … & Mann, M. (2018). Online parallel accumulation–serial fragmentation (PASEF) with a novel trapped ion mobility mass spectrometer. Molecular & Cellular Proteomics, 17(12), 2534–2545.

19 Andrews, G.L., Simons, B.L., Young, J.B., Hawkridge, A.M. and Muddiman, D.C., 2011. Performance characteristics of a new hybrid quadrupole time-of-flight tandem mass spectrometer (TripleTOF 5600). Analytical chemistry, 83(13), pp.5442–5446.

20 Chernushevich, I. V., Loboda, A. V., & Thomson, B. A. (2001). An introduction to quadrupole–time-of-flight mass spectrometry. Journal of mass spectrometry, 36(8), 849–865.

21 Chernushevich, I. V., Merenbloom, S. I., Liu, S., & Bloomfield, N. (2017). A W-Geometry ortho-TOF MS with high resolution and up to 100% duty cycle for MS/MS. Journal of The American Society for Mass Spectrometry, 28(10), 2143–2150.

22 Andrien, B. A., Whitehouse, C., & Sansone, M. A. (1998). Proceedings of the 46th ASMS Conference on Mass Spectrometry and Allied Topics. May 31–June 4, Orlando, FL, 889–890.

23 Franzen, J., (1998). Method and device for orthogonal ion injection into a time-of-flight mass spectrometer. United States Patent US5763878A.

24 Hardman, M., & Makarov, A. A. (2003). Interfacing the orbitrap mass analyzer to an electrospray ion source. Analytical chemistry, 75(7), 1699–1705.

25 Brais, C. J., Ibañez, J. O., Schwartz, A. J., & Ray, S. J. (2021). Recent advances in instrumental approaches to time-of-flight mass spectrometry. Mass Spectrometry Reviews, 40(5), 647–669.

26 Grinfeld, D. and Makarov, A., (2013). Multi-reflection mass spectrometer, United States Patent US9136102B2.

27 Stewart, H., Grinfeld, D., and Makarov, A., (2019). Multi-reflection mass spectrometer, United States Patent US10964520B2.

28 Hauschild, J. P., Peterson, A. C., Couzijn, E., Denisov, E., Chernyshev, D., Hock, C., … & Makarov, A. (2020). A Novel Family of Quadrupole-Orbitrap Mass Spectrometers for a Broad Range of Analytical Applications. Preprints, 2020060111.

29 Michalski, A., Damoc, E., Lange, O., Denisov, E., Nolting, D., Müller, M., … & Makarov, A. (2012). Ultra high resolution linear ion trap Orbitrap mass spectrometer (Orbitrap Elite) facilitates top down LC MS/MS and versatile peptide fragmentation modes. Molecular & Cellular Proteomics, 11(3).

30 Lange, O.; Damoc, E.; Wieghaus, A.; Makarov, A. Enhanced FT for Orbitrap Mass Spectrometry. Proceedings of the 59th ASMS Conference on Mass Spectrometry and Allied Topics, Denver, CO, 2011.

31 Lange, O., Damoc, E., Wieghaus, A., & Makarov, A. (2014). Enhanced Fourier transform for Orbitrap mass spectrometry. International Journal of Mass Spectrometry, 369, 16–22.

32 Douglas, D. J., & French, J. B. (1992). Collisional focusing effects in radio frequency quadrupoles. Journal of the American Society for Mass Spectrometry, 3(4), 398–408.

33 Stewart, H., Hock, C., Giannakopulos, A., Grinfeld, D., Heming, R. and Makarov, A., A rectilinear pulsed-extraction ion trap with auxiliary axial DC trapping electrodes, Proceedings of the 66th ASMS Conference on Mass Spectrometry and Allied Topics, San Antonio, TX, 2018.

34 Sudakov, M., & Kumashiro, S. (2011). TOF systems with two-directional isochronous motion. Nuclear Instruments and Methods in Physics Research Section A: Accelerators, Spectrometers, Detectors and Associated Equipment, 645(1), 210–215.

35 Verentchikov, A. N., Yavor, M. I., Hasin, Y. I., & Gavrik, M. A. (2005). Multireflection planar time-of-flight mass analyzer. I: An analyzer for a parallel tandem spectrometer. Technical Physics, 50(1), 73–81.

36 Nazarenko, L.M., Sekunova L.M., Yakushev E.M., (1989) Time-of-flight mass spectrometer with multiple reflections, Soviet Patent No. SU1725289.

37 Schwartz, J. C., Xaio-Guang, Z. & Bier, M. E. (1995). Method and apparatus of increasing dynamic range and sensitivity of a mass spectrometer, United States Patent No. US5572022A.

38 Denisov, E., Damoc, E., & Makarov, A. (2021). Exploring frontiers of orbitrap performance for long transients. International Journal of Mass Spectrometry, 466, 116607.

39 Rose, R. J., Damoc, E., Denisov, E., Makarov, A., & Heck, A. J. (2012). High-sensitivity Orbitrap mass analysis of intact macromolecular assemblies. Nature methods, 9(11), 1084–1086.

40 Liu, R., Li, Q., & Smith, L. M. (2014). Detection of large ions in time-of-flight mass spectrometry: effects of ion mass and acceleration voltage on microchannel plate detector response. Journal of The American Society for Mass Spectrometry, 25(8), 1374–1383.

41 Blume, J. E., Manning, W. C., Troiano, G., Hornburg, D., Figa, M., Hesterberg, L., … & Farokhzad, O. C. (2020). Rapid, deep and precise profiling of the plasma proteome with multi-nanoparticle protein corona. Nature Communications, 11(1), 3662.

42 Li, J., Cai, Z., Bomgarden, R. D., Pike, I., Kuhn, K., Rogers, J. C., … & Paulo, J. A. (2021). TMTpro-18plex: the expanded and complete set of TMTpro reagents for sample multiplexing. Journal of proteome research, 20(5), 2964–2972.

