## Supplemental Data for "Parallelized Acquisition of Orbitrap and Astral Analyzers Enables High-Throughput Quantitative Analysis"

### Supplementary Data

**Analyzer Mechanics and Vacuum system**: Figure S1 shows a top-down photograph of the Astral analyzer within its vacuum chamber, with the top lid open. The main elements are annotated. An inner lid covering the UHV region (ion mirrors, ion foil and detector) was also removed. Differential pumping was applied by a single multi-stage pump (Pfeiffer Vacuum), the same type used in the instrument's front end and the Orbitrap Exploris 480 design which served as its base[28]. This pumping is made over five separate regions from the ion processor, its surrounding sub-chamber, the injection optics, an additional region at the end of the injection optics, and the UHV region. A sixth region around the transfer multipole was evacuated by an additional small 67 l/s turbomolecular pump. Rough pumping was made by a single two-inlet nXL110iDE dry pump (Edwards Vacuum).

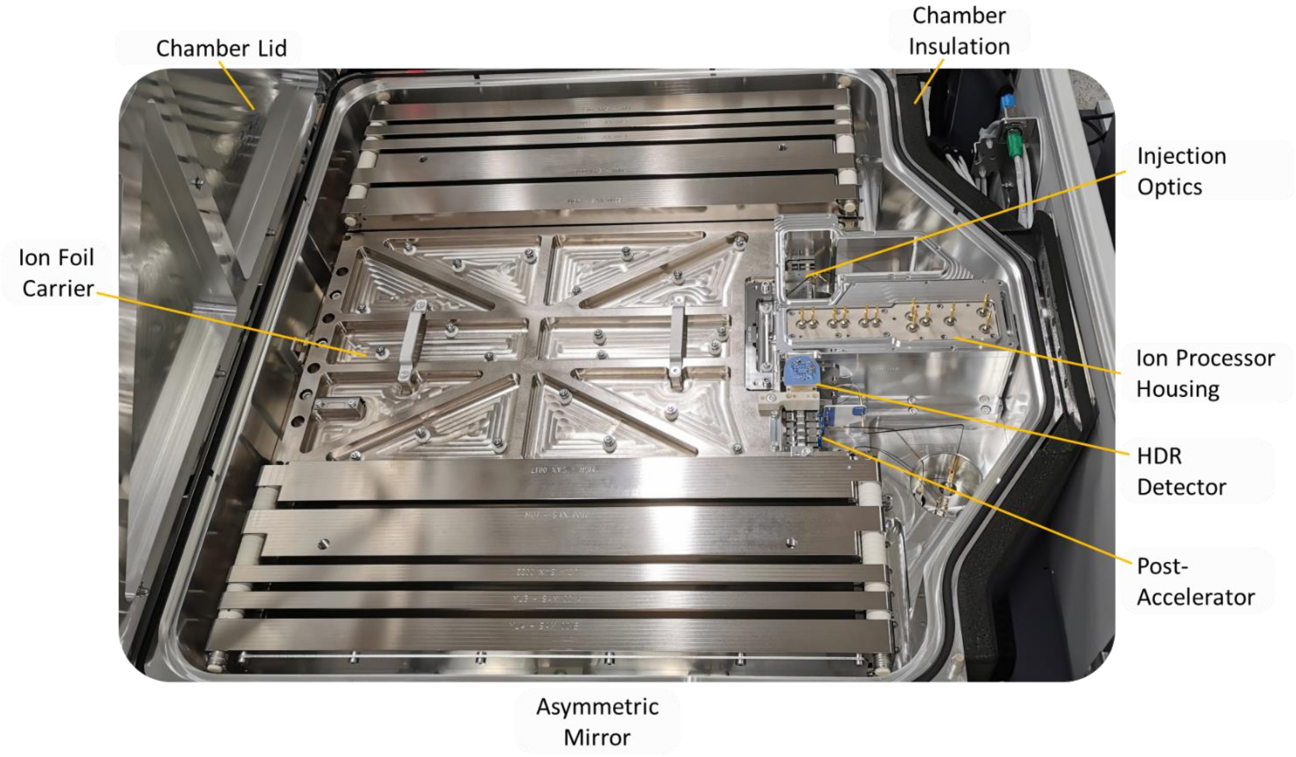


*Figure S1. Photograph of the Astral analyzer*

**Dynamic Range**: The digitizer covers a range from the ~4mV noise threshold to just under 5V, implying a >1000:1 linear dynamic range. Figure S2a and b show scans of inject time for Flexmix ions at m/z 142, 524 and 2122 plotted against the number of ions detected, as directly calculated from the peak area and the calibrated single ion response of the detector. The apparent linear response appears far greater than might be assumed from the detector's range, especially for the two higher m/z which appear linear to closer to 4-orders of dynamic range from a single ion, than 3-orders.

Due to the 30m+ flight path, most ion packets are much wider than the detector's ~1.7ns single ion response, so many more ions are normally required to saturate the detector. Furthermore, as Figure S2c shows for m/z 524 MRFA ions, peak resolution drops with increasing number of ions in-peak due to space charge, an effect that is interestingly weaker when other Flexmix ions are present in a full-MS. This is likely due to space charge in the trap increasing the ion packet size and consequently reducing the charge density of ions in flight. Peak width increases under space charge, further reducing the probability or proportion of detector saturation. Lower m/z saturates earlier as peaks are narrower in time, but even m/z 142 has a reasonably linear response to several thousand ions. This effect is likely an important factor in the instrument's strong performance in label-free quantitation. Besides detector performance, linearity is expected to be affected by space charge effects limiting the capacity of the ion processor, which has a relatively narrow 2mm radius channel, and likely also broadening of the ion packet in flight impacting transmission.

Unlike FTMS analyzers, the presence of intense peaks does not raise the noise baseline throughout the spectrum, aside from a short period of pre-amplifier recovery. Figure S2d shows the isolated MRFA isotopic envelope recorded in a single shot under massive ion load resulting in 40K detected ions (without correction for non-linearity). The measured m/z was not greatly perturbed by the overload. Inset is shown the same mass spectrum but zoomed to show extremely low-lying peaks, even down below the 0.01% level indicative of single ion impacts. The dead zones after the main intense peaks are due to these peaks' own substantial width under space charge, and the time required for the pre-amplifier to recover.

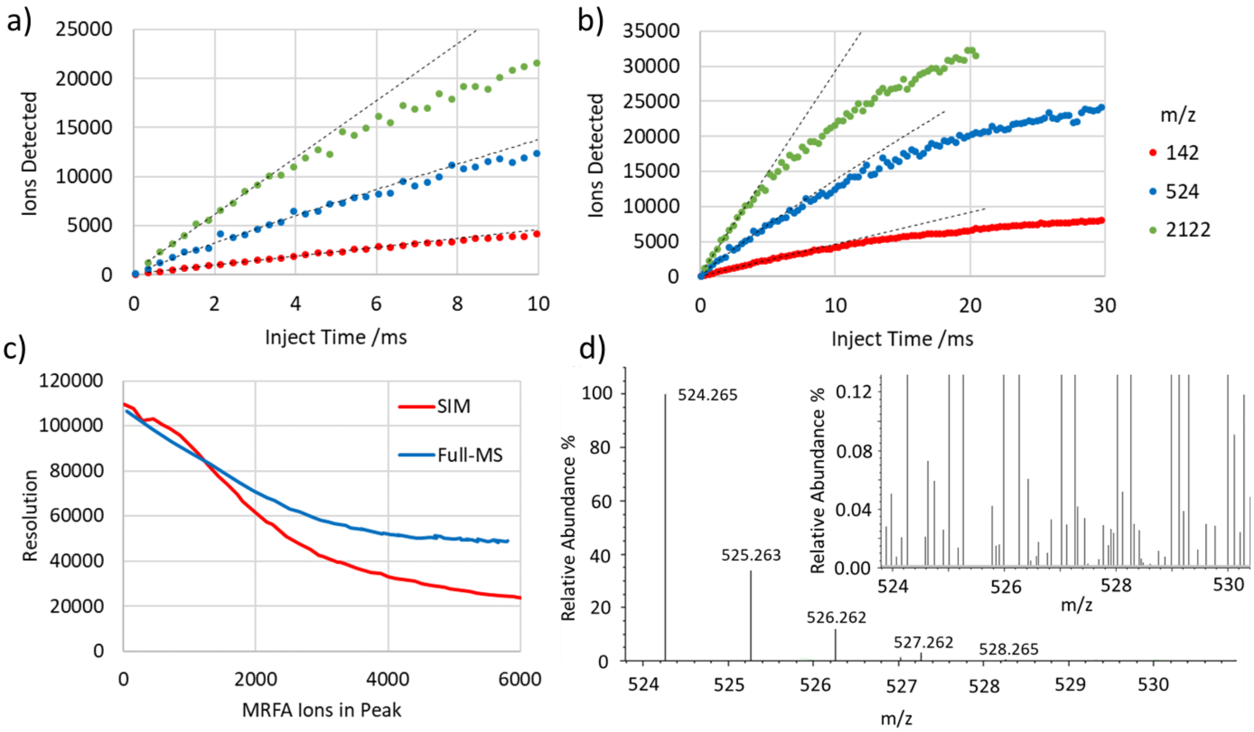


*Figure S2. a) and b) Differing zoom levels of inject time scans of three isolated m/z ions, showing m/z dependent loss of linearity at high ion number. c) Change in resolution of the m/z 524 peak under increasing in-peak space charge, both isolated SIM and in FlexMix Full-MS. d) MRFA isotopic envelope with overloaded 40K detected ions, zoomed inset shows small peaks, including those corresponding to single ions (0.01% level).*

**Summary of Methods:**

Table S1 below shows a summary of the experimental methods used to characterize the Orbitrap Astral mass spectrometer, for reference. In all cases a Vanquish Neo UHPLC system was used to provide chromatographic separation. Further application results that added excess material to the main text are also provided below.

| Method | Sample and Series | Column | Cycle Length | Special | Orbitrap | Astral | Processing |
|---|---|---|---|---|---|---|---|
| **HeLa DIA Throughput and Depth** | 300, 180, 100, & 60, SPD, 200 ng Pierce HeLa | Easy-Spray PepMap 150µm x 15cm | 4.8, 8, 14.4 & 24-min | Trap and elute | R:240k m/z: 380-900 AGC: 300% | Iso: 2 Th Inject: 3ms m/z: 150-2k AGC: 500% | Proteome Discoverer 3.1 software with CHIMERYS |
| | 30 SPD, 1000 ng HeLa | µPAC Neo 110cm | 48-min | Trap and elute | | | |
| **Label-Free Quantitation** | HeLa, E. coli & Yeast 3-Proteome Mix 2:3:1 & 1:3:2ratios, 500 ng | µPAC Neo 50cm | 24-min | Direct injection | R:240k m/z: 380-900 AGC: 100% | Iso: 2 Th Inject: 3 ms m/z: 150-2k AGC: 500% | Spectronaut 17, directDIA |
| **Low Load HeLa** | Pierce Hela 50pg-10ng, 80 SPD | µPAC Neo low load 50cm | 18-min | Direct injection FAIMS -50 V CV | R:240k m/z: 400-800 AGC: 300% | Iso: 20 to 5 Th Inject: 40 to 10 ms m/z: 150-2k AGC: 800% | Spectronaut 17, Direct DIA and Spectral Library Search |
| **Plasma Proteomics** | Neat Ppasma digest 200ng | Easy-Spray PepMap 150µm x 15cm | 8-min | Trap and elute | R:240k m/z: 380-900 AGC: 100% | Iso: 3 Th Inject: 7 ms m/z: 150-2k AGC: 500% | Proteome Discoverer 3.1 software with CHIMERYS |
| | Plasma digest 1000ng | Easy-Spray PepMap Neo 75um x 75cm | 60-min | Direct injection | R:240k m/z: 380-900 AGC: 100% | Iso: 2 Th Inject: 7 ms m/z: 150-2k AGC: 500% | Proteome Discoverer 3.1 software with CHIMERYS |
| **TMT Quantitation** | TMT11plex Yeast Digest 25 to 500ng | Easy-Spray PepMap Neo 75um x 75cm | 70-min gradient | Direct injection FAIMS -40, -60, -80V CV | R:180k m/z: 400-1.5k AGC: 200% | Iso: 0.5 Th Inject: 20 ms m/z: 110-2k | Proteome Discoverer 3.1 software |

| | | | | | | | |
|---|---|---|---|---|---|---|---|
| | | | | | | AGC: 200% NCE:33 | |
| **Large Cohort HeLa QC** | Pierce HeLa 200ng | Easy-Spray PepMap150µm x 15cm | 14.4-min | Trap and elute | R:240k m/z: 380-900 AGC: 100% | Iso: 2 Th Inject: 3 ms m/z: 150-2k AGC: 500% | Proteome Discoverer 3.1 software with CHIMERYS |

*Table S1. Summary of experimental methods.*

**Plasma Proteomics**: The plasma proteome has been particularly difficult to interrogate due to high concentration, dominant proteins such as albumin exhausting the dynamic range of analysis. Deep analysis requires dynamic range compression via techniques such as immunodepletion, chromatographic fractionation or nanoparticle enrichment.[41] Digested neat plasma was measured with 180 SPD/8-min and 24 SPD/60-min methods, separated on 150um x 15cm PepMap and 75um x 75cm Easy-Spray PepMap Neo columns respectively. The isolation window was set to 3 Th for the former test and 2 Th in the latter, with a 7ms max injection time. Data processing was performed in Proteome Discoverer 3.1 with Chimerys and returned over 600 protein identifications in the 8-minute method and more than 1100 in 60 minutes, as shown in Figure S3. Whilst only a fraction of the plasma proteome, this is nevertheless a high analytical depth to achieve at this throughput and without enrichment.

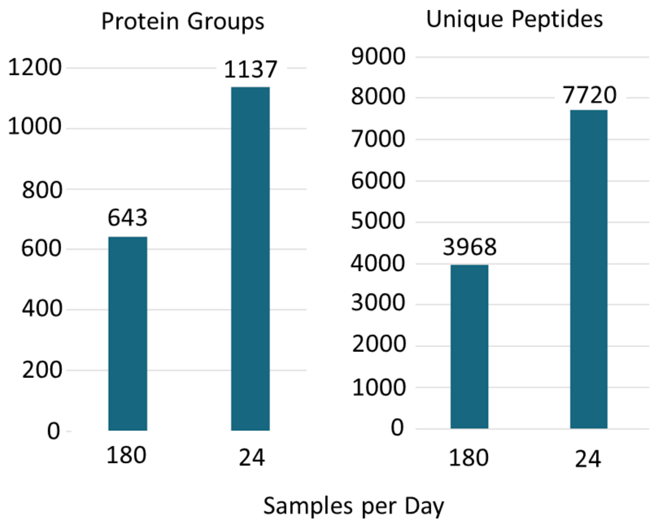


*Figure S3. Protein and peptide ID results from analysis of 200 and 1000ng neat plasma at 180 and 24 samples per day respectively.*

**Tandem Mass Tags**: The Astral analyzer demonstrates sufficient >50K resolution around m/z 130 to separate the reporter ion doublets of highly multiplexed tandem mass tags, such as Pierce TMT11plex$^{TM}$ isobaric labels and up to the current multiplexing limit of 18[42]. Figure S4a shows the peak profile of one such reporter doublet, generated from MS/MS of electrosprayed TMT11plex reagent sample. Notably in this example the high gain channel is saturated by the intense reporter signal, and the low gain channel used instead. Figure S4b shows an example of an MS/MS spectra obtained of an entire range of TMT reporter ions. The peak doublets are well resolved and all present, whilst the average mass deviation was 0.42ppm.

A TMT quantitation dilution series experiment was run with injections of 25, 50, 100, 200 and 500ng Pierce TMT11plex™ yeast digest, a standard for benchmarking TMT methods, separated over 50 minutes on a 75um x 75cm Easy-Spray PepMap Neo UHPLC column. Three FAIMS compensation voltages were used: -40, -60 and -80V. The Full-MS settings used were 180,000 resolution, with a scan range of 400-1500 and an AGC target of 200%. MS/MS settings incorporated a max inject time of 20 ms, a 200% AGC target, a first mass of 110 and normalized collision energy of 35.

After processing in Proteome Discoverer, the reported protein groups and unique peptides are shown in Figure S5 c and d, as well as the interference-free index for Met6 (Figure S4 e)

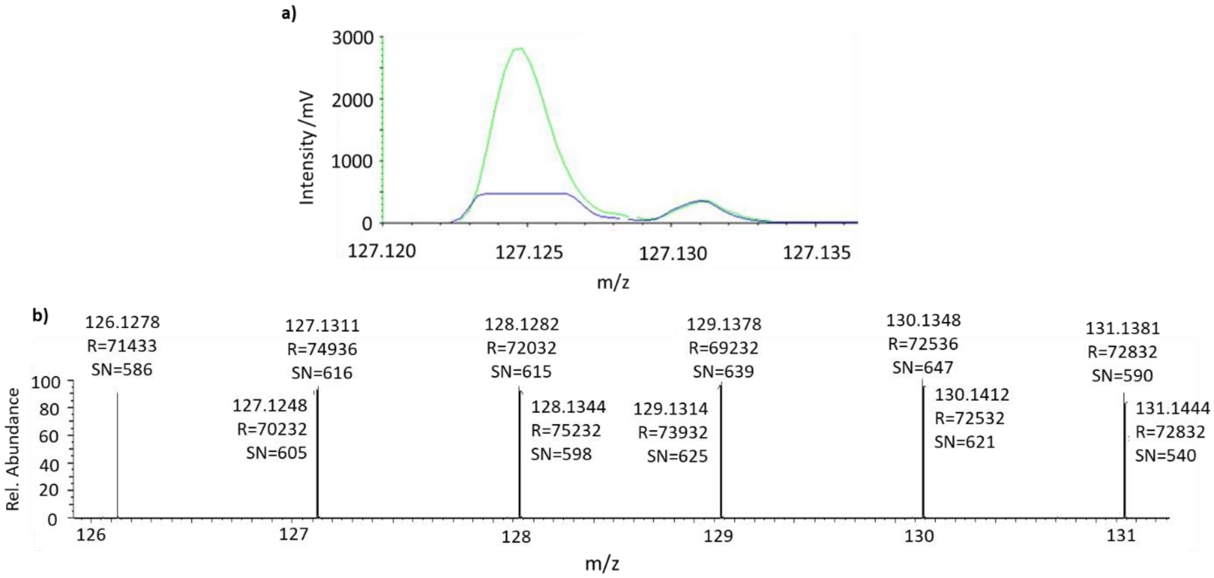


*Figure S4. a) Peak profile of a resolved TMT reporter doublet. b) TMT11plex reporter ion mass spectrum.*

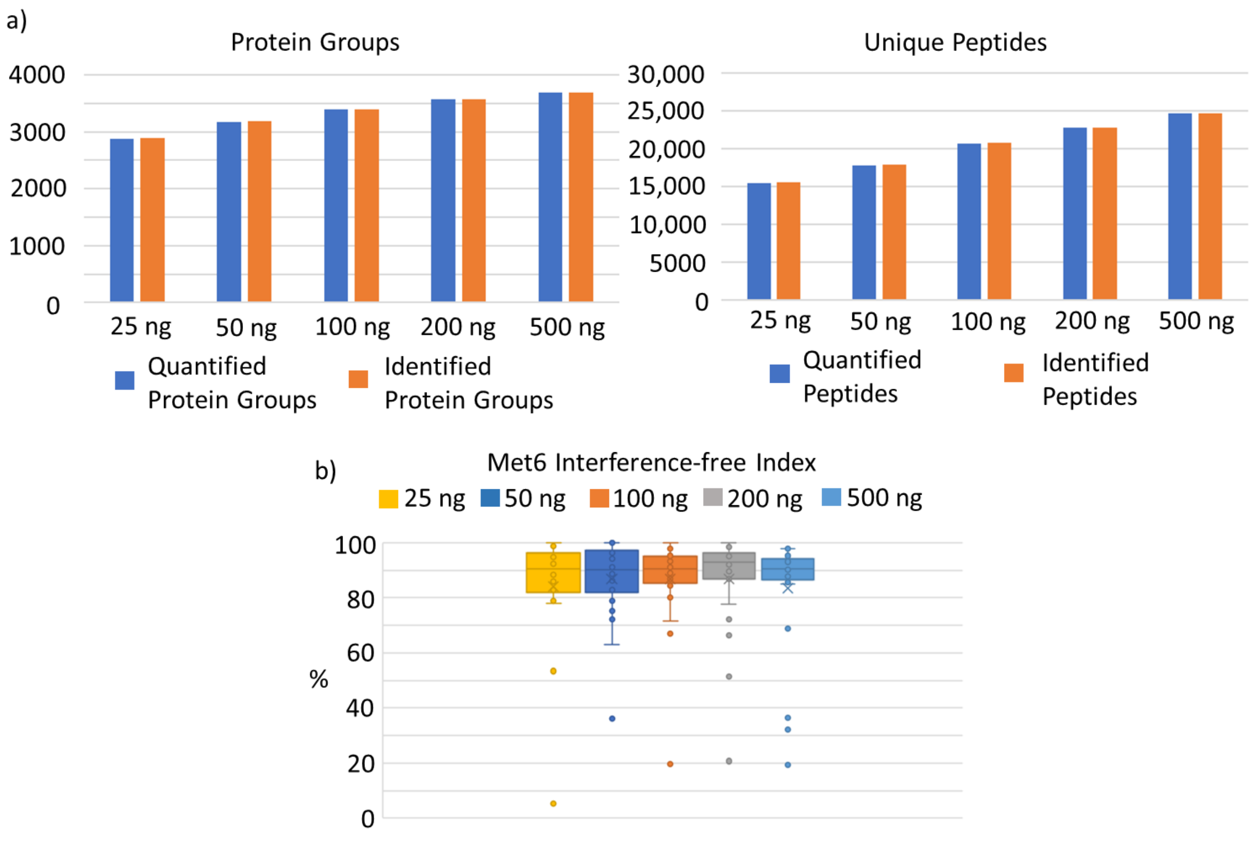


*Figure S5. a) Protein groups and unique peptides identified and quantified from different sample loads. b) Interference-free index (IFI) for Met6 at different sample loads (IFI 100 reflects no interference).*

**Large Cohort Studies**: Many patient cohort studies measure large numbers of samples, with a prerequisite that the mass spectrometer be sufficiently robust and generate reproducible data over the course of the experiment series. To test this, a thousand 100ng neat plasma injections were made through a PepMap 150µm x 15cmcolumn, with the 11-minute gradient 100 samples per day DIA method. Regular 200 ng HeLa injections were made to monitor performance degradation, and the Proteome Discoverer processed results shown in Figure S5. NoI decay in the number of identified peptides or protein groups could be observed, with no cleaning/maintenance of the front-end or recalibration required.

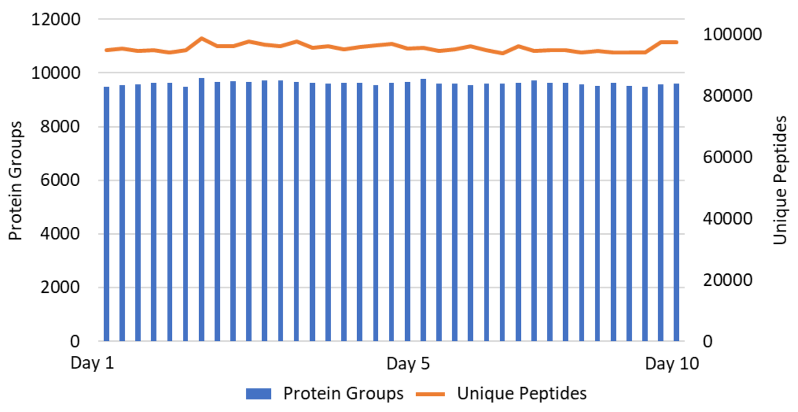


*Figure S5. 200ng, 100 samples per day HeLa injections over 10 days, interspersed within a large cohort study of 1000, 100ng neat plasma injections using the same analysis method.*
